# Adaptive loss of function accelerated the evolution of ancient and modern human cognition

**DOI:** 10.64898/2026.04.27.721126

**Authors:** Alexander L. Starr, Gabriella M. Cale, Leslie Magtanong, Michael E. Palmer, Hunter B. Fraser

## Abstract

Methods to detect accelerated evolution have identified many genomic regions with unexpectedly rapid evolution in the human lineage—significantly more than in chimpanzees, our closest living relatives. However, these methods focus on accelerated sequence evolution of short non-coding regions, leaving open the questions of how to identify accelerated evolution of molecular function, as opposed to sequence, and whether accelerated evolution has shaped the human genome more broadly. Here, we introduce a new approach to detect accelerated evolution: Function-Aware Statistical Test for Evolutionary Rates (FASTER). In contrast to previous methods, FASTER can detect not only accelerated evolution of sequence, but also of predicted function, and can be applied to any set of genomic regions. Applying this method to humans and chimpanzees, we identified protein-coding, untranslated (UTR), and non-coding regions with accelerated evolution of function. Across all these genomic levels, we consistently found more rapid evolution in conserved sites in the human lineage compared to chimpanzee, many of which are predicted to reduce protein stability or chromatin accessibility. Multiple lines of evidence suggest this human-acceleration was driven in part by positive selection on brain development and cognition which has continued to shape human evolution even in the past several thousand years. Collectively, these results demonstrate the power of genome-wide scans for the evolution of predicted function and specifically suggest that a higher rate of reduction in function—including widespread decreases in *cis*-regulatory activity—may have been an important driver of both ancient and recent human evolution.

## Introduction

Understanding the genetic basis of human evolution remains a central goal in evolutionary biology and medical genetics. Both early comparative studies and more recent genome-wide analyses suggest that changes in non-coding *cis*-regulatory elements (CREs) that affect gene expression, rather than protein-coding substitutions, may have played a dominant role in human evolution^1–3^. In recent years, advances in comparative genomics have enabled systematic scans for small genomic regions that exhibit signatures of accelerated evolution, defined as an excess of substitutions along a particular lineage relative to some neutral expectation^4–7^. These approaches have identified hundreds of human accelerated regions (HARs), the majority of which are located in non-coding portions of the genome and have *cis*-regulatory activity^4–7^. Interestingly, some^4,8^ (but not all^6^) studies have reported a greater number of accelerated regions in the human lineage compared to the chimpanzee lineage. Although it is unclear exactly how much of the bias toward accelerated evolution in the human lineage is due to relaxed negative selection, many HARs may be the result of positive selection^9^.

The identification of an excess of accelerated regions in the human lineage raises several questions. First, as outlined above, methods to detect accelerated evolution are typically optimized for individual short non-coding elements^6,7,10,11^. As a result, it is not clear whether these findings extend to other levels of biological organization such as protein-coding exons, untranslated regions (UTRs), and the *cis*-regulatory context of entire genes, or even whether accelerated evolution exists in these broader contexts. Second, it is unclear what the consequences of more rapid evolution of conserved sites in the human lineage might be. In parallel to these studies, a growing body of work has demonstrated the importance of loss of or reductions in function in human evolution. For example, inactivation of the *CMAH* gene in the human lineage is thought to have altered immune responses, pathogen susceptibility, and reproductive biology^12,13^. Although it is unknown whether inactivation of genes such as *CMAH* was evolutionarily advantageous, decreases in function can provide a fitness advantage. For example, we recently showed that decreased expression of genes linked to autism spectrum disorder was adaptive during human evolution^14^. More broadly, decreases in CRE activity underlie key morphological and physiological changes across diverse taxa^15^. Although random mutations in conserved sites are generally expected to decrease CRE function, it is unclear whether observed substitutions in conserved sites, which have passed through the sieve of natural selection, also tend to disrupt CREs. If this were the case, one potential consequence of the bias toward accelerated evolution in the human lineage could be a bias toward decreases in CRE function.

Collectively, this suggests that more flexible analytical frameworks would enable us to identify human-accelerated evolution more broadly and better understand how it shaped uniquely human traits and disease susceptibility. Here, we introduce FASTER, a new methodology designed to detect accelerated evolution of sequence or predicted function in one lineage (e.g., human) relative to another (e.g., chimp) using any type of variant effect prediction as input. Applying our approach in conjunction with conservation scores as a predictor of functional impact, we identified human acceleration in the protein-coding sequences, UTRs, and entire *cis*-regulatory landscapes of genes and pathways. Across all levels of biological organization, we consistently found more rapid evolution in conserved sites in the human lineage than in the chimpanzee lineage. Moreover, incorporating machine learning-derived predictions of the effects of substitutions on chromatin accessibility (CA) into FASTER, we identified genes with accelerated evolution of CA and found that substitutions in more highly conserved sites are predicted to decrease accessibility much more frequently than those in unconserved sites. Finally, we found that even in the past 10,000 years of human evolution, positive selection often occurred in highly conserved sites, likely decreased CA in fetal cortical neurons, and affected cognitive and psychiatric traits. In aggregate, these results demonstrate that human-accelerated evolution is widespread across diverse genomic features, potentially biasing human evolution toward reduced function and suggesting that decreases in *cis*-regulatory activity may have been particularly widespread during both ancient and modern human evolution.

## Results

At their core, most methods to detect accelerated evolution compare the number of substitutions in a given region relative to the number expected under some model of evolution, often based on empirical measurements across a phylogeny^4,6,7^. For example, human ancestor quickly evolved regions (HAQERs) were defined as regions 500 bases in length with at least 29 human-derived genetic differences in that region, giving equal weight to each substitution^16^. Similarly, FASTER starts by computing the number of substitutions separately in two sister lineages (in this case humans and chimpanzees). However, rather than treating all substitutions equally, a key feature of our approach is that it can flexibly give more weight to those more likely to affect phenotype and fitness. In this work, we weight each substitution by its evolutionary conservation or its predicted effect on CA to account for its likely functional consequences (Fig. 1A).

**Figure 1:**
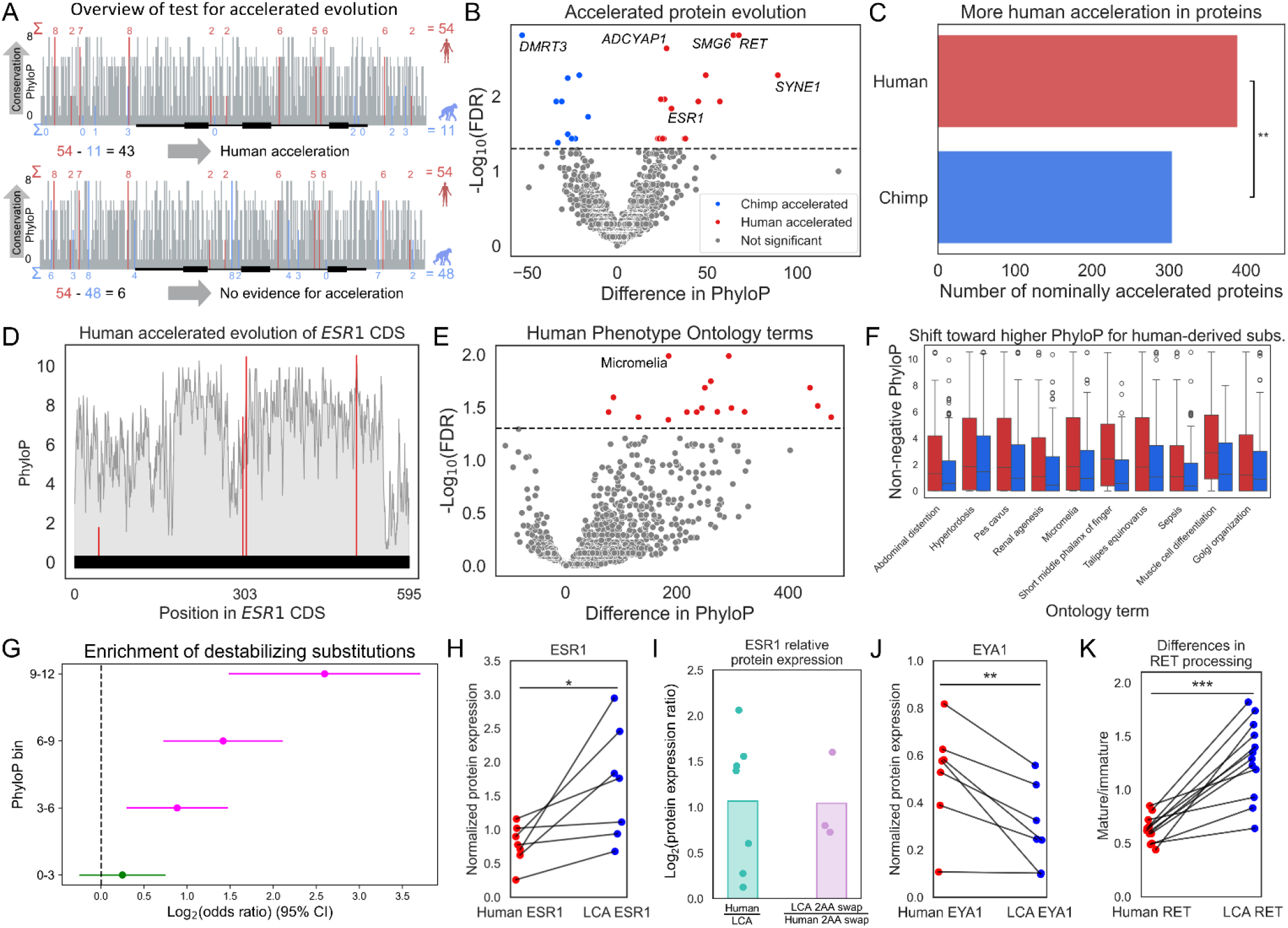
Identification and characterization of human accelerated proteins (HAPs). **A)** Overview of the FASTER method to identify accelerated evolution. A hypothetical accelerated example is depicted on top and non-accelerated example on bottom. Red and blue numbers indicate PhyloP scores for sites with human-derived and chimpanzee-derived substitutions, respectively, with their sums shown on the right. **B)** PhyloP differences driving protein acceleration. **C)** Nominally human-accelerated and chimpanzee-accelerated proteins. ** p < 0.005. **D)** PhyloP scores in the *ESR1* CDS. Position along the x-axis is the position in the amino acid sequence in the canonical ESR1 protein. Red indicates sites with human-derived substitutions and grey indicates sites with no substitutions. **E)** PhyloP differences driving acceleration of Human Phenotype Onotology (HPO) terms. **F)** PhyloP distributions for human-derived (red) and chimpanzee-derived (blue) nonsynonymous substitutions for proteins in each of the ten significant HPO or Gene Ontology Biological Process (GOBP) categories. **G)** Enrichment of predicted destabilizing substitutions in conserved sites. **H)** Normalized protein expression for human and human-chimpanzee last common ancestor (LCA) ESR1. Lines indicate paired measurements from paired biological replicates on the same Western blot. * p < 0.05. **I)** Log_2_ fold-change for normalized human ESR1 protein expression divided by human-chimpanzee LCA expression (left, computed from the data in H) compared to swapping only the amino acids at positions 300 and 306 (right); the only difference between the two comparisons is in the three human-derived amino acid changes outside of positions 300 and 306, which have no measurable effect. **J)** Normalized protein expression for human and human-chimpanzee LCA EYA1. ** p < 0.005. **K)** Ratio of mature, glycosylated RET expression to immature, unglycosylated RET expression. *** p < 0.0005.

Identifying accelerated evolution necessarily involves a comparison, such as to an expected non-accelerated rate. However, this expectation varies drastically based on factors such as the local mutation rate, density of functionally important sites, etc. Rather than try to compute this expectation, we instead compared orthologous regions in two sister lineages and used the difference in the sums of predicted effects of lineage-specific substitutions (defined here as sites in which the human allele differs from the chimpanzee allele and one of the alleles matches the gorilla allele; see Methods) as our test statistic. Throughout, we refer to significantly more rapid evolution in one lineage compared to the other as “acceleration” on the basis that, in most cases, evolution in the accelerated lineage is more rapid relative to gorilla as well (Supp. Text 1).

As a simple example, consider a region with four human-derived substitutions, each with a predicted-effect weight of 5, and four chimpanzee-derived substitutions each with a weight of 1. The weighted sums are therefore 20 for human and 4 for chimpanzee, yielding a test statistic of 16 despite identical substitution counts. To determine whether this difference is greater than expected by chance, FASTER repeatedly reassigns the observed substitutions between lineages and genomic regions according to their genome-wide background probabilities and recalculates the test statistic; the observed value is then compared with this null distribution to obtain a p-value (see Methods for details). Thus, unlike approaches such as dN/dS, which compare rates based on numbers of substitutions and treat substitutions within a class equivalently, FASTER can detect acceleration arising either from a greater number of substitutions (“count-driven” acceleration) or from substitutions with systematically larger predicted functional effects (“weight-driven” acceleration), or both.

### Human-accelerated evolution in protein-coding genes likely decreased protein stability

As an initial application of FASTER, we applied it to nonsynonymous substitutions (those that affect amino acid sequence), weighting substitutions by their PhyloP scores, a metric of evolutionary conservation often used as a proxy for functional importance^8,11^, while controlling for global interspecies difference in the substitution rate of each amino acid. Importantly, these PhyloP scores were computed without information from the human, chimpanzee, or bonobo genomes, ensuring they are independent of human- and chimpanzee-derived substitutions. We identified 16 human-accelerated proteins (HAPs) and 10 chimpanzee-accelerated proteins (CAPs) at a false discovery rate (FDR) of 0.05 (Fig. 1B, Supp. Table 1). Notably, there were significantly more nominally significant (unadjusted p-value < 0.05) HAPs than CAPs (Fig. 1C, 1.3-fold more nominal HAPs, p = 0.0014, binomial test; this increases to 1.4-fold more nominal HAPs with a more strict definition of acceleration, Supp. Text 1), suggesting that the previously observed enrichment of human-acceleration in conserved non-coding elements (such as most HARs) extends to protein-coding regions (see Supp. Text 2 for comparison to previous results derived from dN/dS-like approaches).

The most strongly human-accelerated protein was the oncogene RET, a tyrosine kinase required for the development of several organs—including neuronal proliferation, survival, and migration in the brain and peripheral nervous system—that is often mutated in thyroid cancer^17^. Of its 10 nonsynonymous substitutions in the human lineage, three were in sites that are conserved in all 446 other placental mammals in our study; in contrast, RET had zero nonsynonymous substitutions in chimpanzee (Supp. Fig. 1A). Another oncogene and important regulator of cognitive function, the estrogen-activated transcription factor ESR^18^, had 5 nonsynonymous substitutions (2 conserved in all 446 placental mammals) compared to zero in chimpanzee (Fig. 1D). Interestingly, 2 of the substitutions were within 3 amino acids of K303, somatic mutations in which can cause breast cancer by affecting the activity and degradation of ESR^19^.

FASTER’s flexibility allows for any genomic regions as input, including any gene set of interest. Therefore, we next tested for accelerated coding sequence evolution among groups of proteins associated with particular human phenotypes (using the Human Phenotype Ontology^20^ (HPO)) or biological processes (Gene Ontology Biological Process^21^ (GOBP)). We identified 17 HPO terms (Fig. 1E) and 5 GOBP terms (Supp. Fig. 1B-C) with accelerated protein-coding sequence evolution (FDR < 0.05, Supp. Table 2). Notably, all 22 of these were accelerated in humans rather than in chimpanzees. However, some proteins (primarily RET) were such strong outliers that they could dominate enrichment results for small gene sets. To reduce the impact of outliers, we focused on terms that remained nominally significant after removing the three most human-accelerated and three most chimpanzee-accelerated proteins, resulting in 8 HPO and 2 GOBP terms; all 10 were human-accelerated.

Next, we tested whether these human-accelerated gene sets were primarily driven by an excess of either 1) the number of human-derived substitutions (count-driven), or 2) human-derived substitutions occurring at more conserved positions (weight-driven), which are not mutually exclusive (see Methods). We found that all ten gene sets were weight-driven as the human-derived substitutions had significantly greater conservation (PhyloP scores Mann-Whitney U test FDR < 0.05, Fig. 1F). In contrast, only 6/10 gene sets were also count-driven (i.e. had nominally significantly more human-derived substitutions, binomial p < 0.05), emphasizing the impact of weighting by variant effect prediction when testing for accelerated evolution. For example, the human-acceleration of proteins implicated in micromelia (congenital shortness of limbs; chimpanzees notably have shorter legs than humans) was entirely weight-driven; human-derived substitutions occurred at more conserved positions (p = 0.00036, Mann-Whitney U test), with no difference in the number of substitutions (p = 0.40, binomial test). Altogether, these results point to a more dominant role of human-derived substitutions at conserved sites, rather than simply the number of substitutions, driving the strong trend of human-acceleration among protein-coding gene sets.

As discussed above, random mutations in conserved sites are expected to decrease function on average. For example, many disease-causing mutations disrupt protein stability or processing^22^. However, it is unclear whether human-derived fixed substitutions in conserved sites, which are unlikely to be highly deleterious, are also biased toward decreasing protein stability. To investigate this question, we used predictions of the effects of amino acid changes on protein stability (the model used to make these predictions does not take evolutionary conservation into account, preventing any technical circularity in our results^23^). We found that substitutions in the most conserved sites were six times more likely to be predicted to alter stability than those in non-conserved sites (Fig. 1G), suggesting that a major mechanism of action for human-derived substitutions in conserved sites is to decrease protein stability, which may reduce protein function.

We next asked whether human-derived nonsynonymous substitutions at conserved sites might alter protein abundance or processing. We selected three human-accelerated and two nominally human-accelerated proteins (for a total of five, selected without regard for their predicted effects on stability; see Methods) and tested for effects of the human-derived substitutions on protein expression. Specifically, we introduced plasmids encoding the human and human-chimpanzee last common ancestor (LCA) coding regions into one of two human cell lines (293T kidney cells for all proteins except ESR1 which was tested in ESR1-negative breast cancer cells) and measured protein levels by Western blot. Of these, 3 out of 5 had significant differences in protein expression or processing (Supp. Fig. 2). First, human ESR1 protein was expressed ∼2-fold lower than LCA ESR1 (p = 0.024, paired t-test, Fig. 1H). Given the proposed role of K303 in regulating the degradation of ESR^19^, we investigated whether the human-derived substitutions in the sites near K303 might be at least partially causal for the difference in protein levels. We found that swapping just those two flanking sites between LCA and human ESR1 was sufficient to recapitulate the 2-fold difference that we observed between the original human and LCA proteins (Fig. 1I), suggesting that the substitutions in these two sites near a breast cancer driver mutation are likely the primary cause of the difference in ESR1 protein levels (although we cannot rule out contributions from other substitutions).

**Figure 2:**
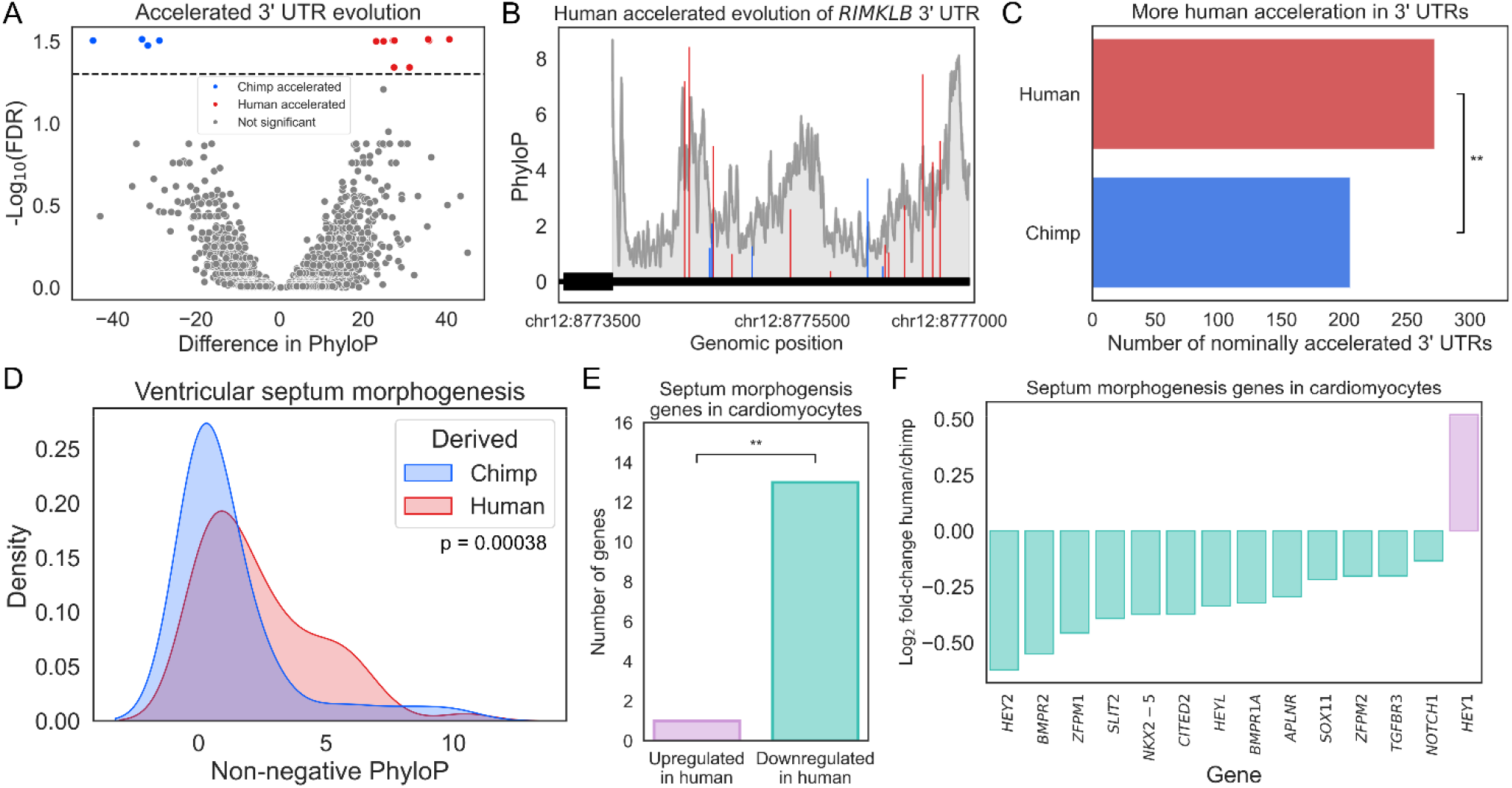
Accelerated evolution of UTRs in human and chimpanzee. **A)** 3’ UTR acceleration. Human-accelerated 3’ UTRs are shown in red and chimpanzee-accelerated in blue. **B)** Plot showing PhyloP scores in the *RIMKLB* 3’ UTR. Red indicates sites with human-derived substitutions, blue indicates sites with chimpanzee-derived substitutions. The grey trace is the moving average of PhyloP scores across all sites. **C)** Nominally human-accelerated and chimpanzee-accelerated 3’ UTRs. ** p < 0.005. **D)** PhyloP distributions for human-derived (red) and chimpanzee-derived (blue) substitutions in the 5’ UTRs of genes involved in ventricular septum morphogenesis. **E)** Genes with increased expression (purple) and decreased expression (teal) from the human allele in iPSC-derived human-chimpanzee hybrid cardiomyocytes. ** p < 0.005. **F)** Log_2_ fold-changes of ventricular septum morphogenesis genes from (E). All have FDR < 0.05.

In contrast, we observed different effects of human-accelerated substitutions in two other essential developmental factors, EYA1 and RET. Human EYA1 was more highly expressed than LCA EYA1 (p = 0.0079, paired t-test, Fig. 1J), suggesting increased protein production and/or stability. For RET, we did not observe a significant difference in total expression level (p = 0.75, paired t-test, Supp. Fig. 1D). However, the mature, functional form of RET is extensively glycosylated and is distinguishable from immature RET as it has a higher molecular weight^24^. Despite the lack of difference in total expression, we found that the human RET construct generally produced more immature RET relative to mature RET, whereas the opposite was true for LCA RET (p = 3.6x10^-5^, paired t-test, Fig. 1K). This is consistent with human RET having altered processing relative to LCA RET, although we cannot rule out faster degradation specifically of mature human RET. Collectively, these results suggest that human-derived substitutions in conserved coding sites tend to decrease predicted protein stability, though their effects on individual proteins are highly varied.

### Human-accelerated evolution of UTRs

Comparing PhyloP scores of human vs. chimpanzee-derived substitutions genome-wide, we observed an overall shift towards substitutions occurring in more conserved sites in the human lineage (Supp. Fig. 3). While this pattern could have important implications for human evolution, its global nature does not readily implicate specific types of genes, pathways, traits, or evolutionary processes (such as positive selection vs relaxed constraint). Therefore, in all analyses below we controlled for this global shift, which makes any results of human-acceleration conservative. To do this, we structured the permutations such that the genome-wide shift toward higher PhyloP scores is reflected in the probability that a substitution is assigned to the human or chimpanzee lineage (see Supp. Text 3 for elaboration on the reasoning behind this and Methods for implementation details). This ensures that our null model reflects any genome-wide, lineage-specific shifts.

**Figure 3:**
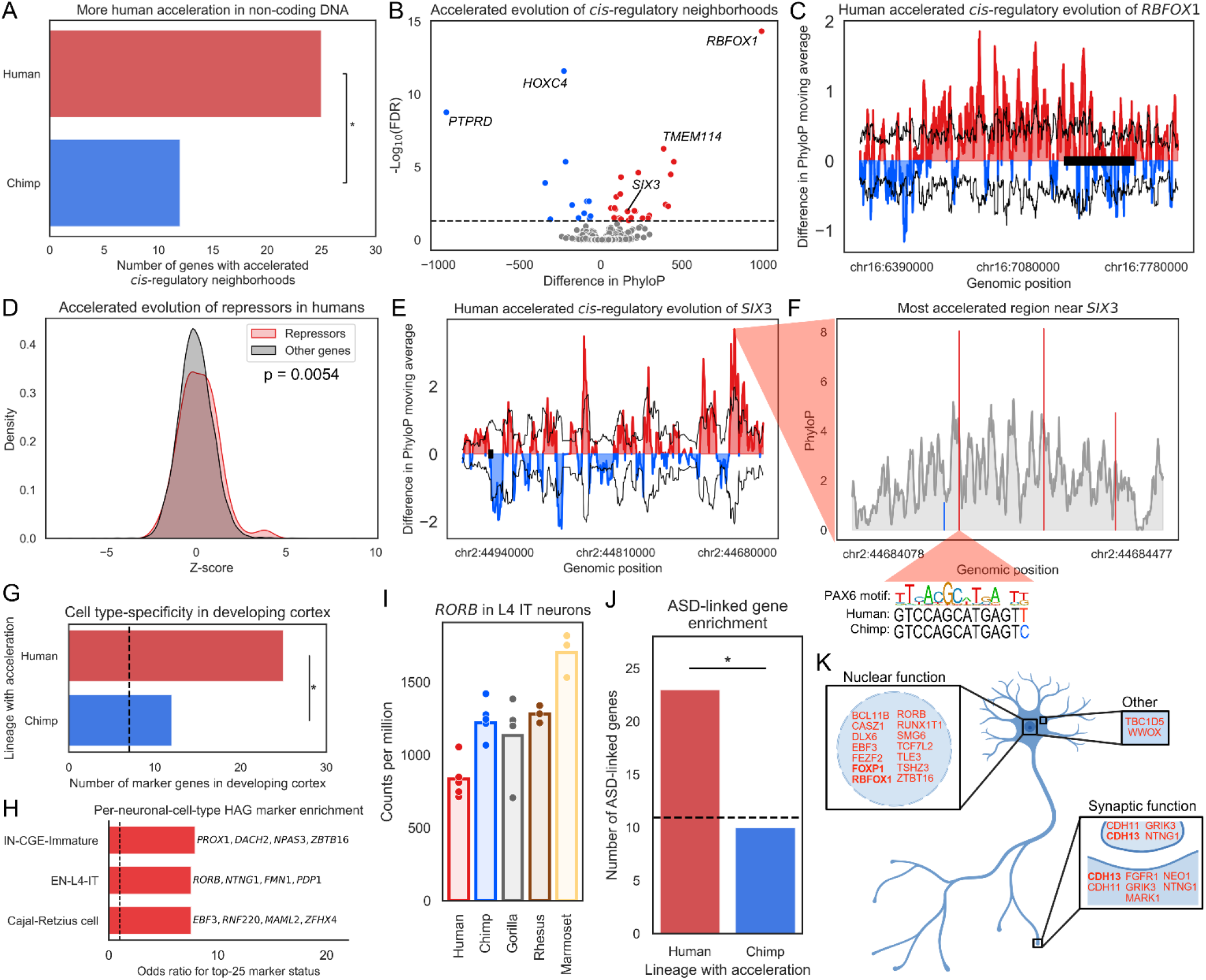
Accelerated evolution of *cis*-regulation in human and chimpanzee. **A)** Number of human-accelerated and chimpanzee-accelerated *cis*-regulatory neighborhoods. * p < 0.05. **B)** Gene-level *cis*-regulatory neighborhood acceleration. HAGs are shown in red and CAGs in blue. **C)** Accelerated evolution of the *RBFOX1 cis*-regulatory landscape in humans. Colored, filled trace shows the moving average of the difference in the sum of human-derived and chimpanzee-derived PhyloP scores. The black trace indicates one standard deviation above and below the mean PhyloP difference of 100 random permutations of which lineage each substitution was assigned to. The black rectangle at y = 0 indicates the *RBFOX1* gene body. **D)** Distribution of z-scores for accelerated evolution for repressors (red) vs. non-TF genes (grey). P-value is from a t-test comparing repressors against allother non-TFs. Corresponding p = 0.80 for activators. **E)** Same as in (C), but for *SIX3* with genomic position inverted. The red cone leading to (F) shows the region that (F) zooms in to. **F)** Plot showing PhyloP scores in the most accelerated non-coding region near *SIX3*. Red indicates sites with human-derived substitutions, blue sites with chimpanzee-derived substitutions. The grey trace is the moving average of PhyloP scores across all sites. At the bottom, the binding motif for PAX6 and corresponding sequences at one human-derived substitution are shown. **G)** Number of nominal HAGs and CAGs that are cell type-specific marker genes in the developing neocortex * p < 0.05. The dashed line represents the number expected based on genes without acceleration. **H)** Nominal HAGs enriched in three neuronal cell types (FDR < 0.05 for each). **I)** Expression of *RORB* in L4 IT neurons. **J)** Number of nominal HAGs and CAGs that are linked to ASD * p < 0.05. The dashed line represents the number expected based on genes without acceleration. **K)** Schematic of a neuron with the subcellular localization of ASD-linked nominal HAGs. HAGs with FDR < 0.05 are bolded.

Applying FASTER to the UTRs of all protein-coding genes, we identified about 2-fold more genes with human-accelerated UTRs (HAUs) than with chimpanzee-accelerated UTRs (3’ UTRs: 9 human-accelerated vs. 4 chimpanzee-accelerated, Fig. 2A-B; 5’ UTRs: 5 human-accelerated vs. 2 chimpanzee-accelerated, Supp Fig. 4A, Supp. Table 3). This difference became more significant as the statistical cutoff was relaxed to allow for a more comprehensive comparison of human- vs chimpanzee-acceleration (3’ UTRs: 1.8-fold more nominal HAUs, p = 0.0025, Fig. 2C; 5’ UTRs: 1.6-fold more nominal HAUs, p = 0.037, binomial test, Supp. Fig. 4B; 2.0-fold and 1.6-fold more nominal 3’ and 5’ HAUs with a more strict definition of acceleration, Supp. Text 1). This suggests that substitutions in conserved UTR sites tend to co-occur in specific genes more often in the human lineage than the chimpanzee lineage, in addition to the global shift toward substitutions in higher PhyloP sites in humans (Supp. Fig. 3). Grouping UTRs by HPO and GOBP terms, we identified acceleration for 2 GOBP terms in 5’ UTRs and none for 3’ UTRs (Supp. Fig. 4C, Supp. Table 4). The most accelerated term was ventricular septum morphogenesis, which is a key process during embryonic heart development^25^. As with the nonsynonymous accelerated gene sets, this was primarily weight-driven as there was a shift toward substitutions in more conserved sites in the human lineage (Fig. 2D, p = 0.00038, Mann-Whitney U test).

**Figure 4:**
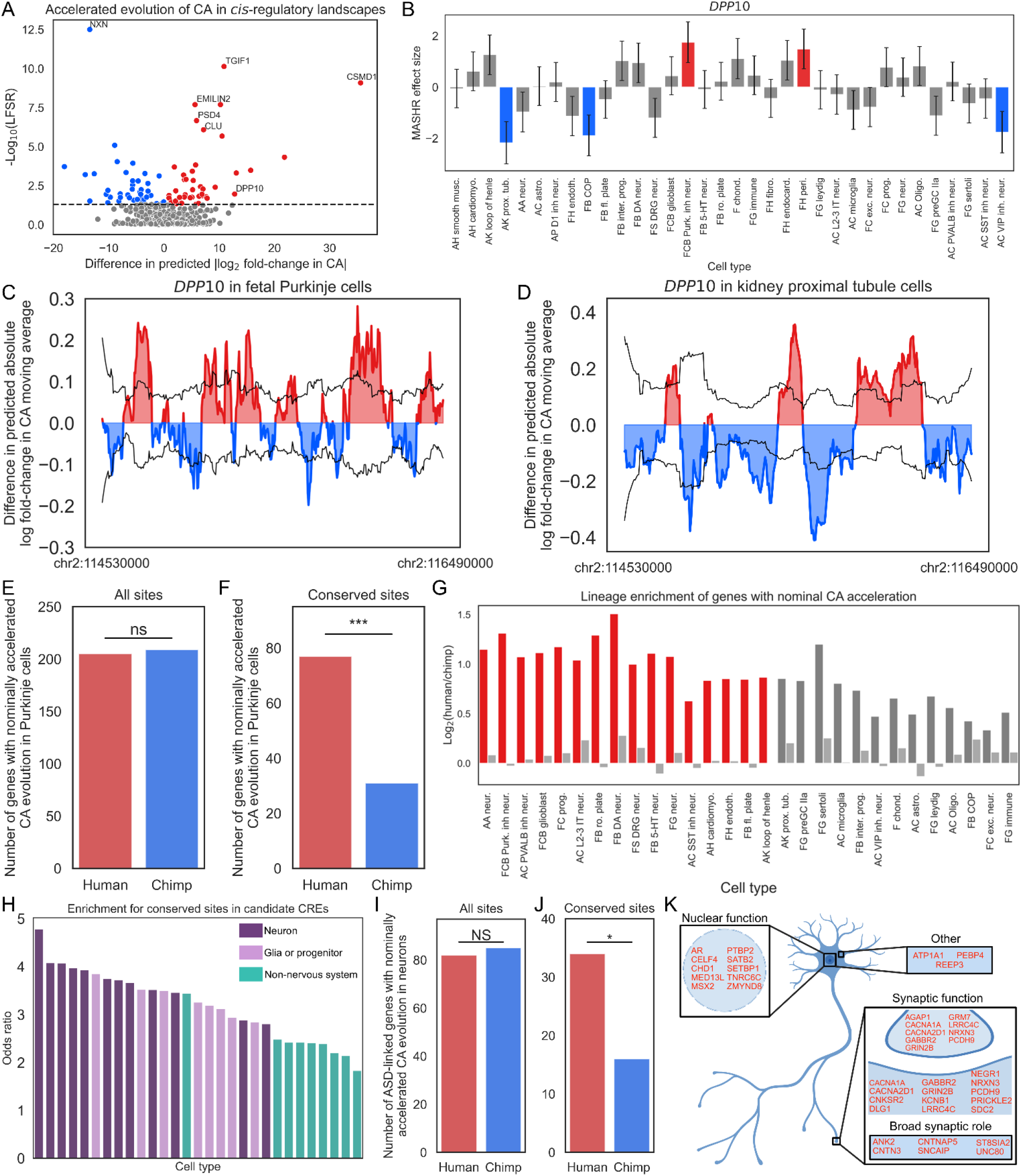
Accelerated evolution of chromatin accessibility in human and chimpanzee. **A)** Genes with accelerated evolution of CA. Human-accelerated CA shown in red and chimpanzee-accelerated CA in blue. **B)** Estimated effect size from mashr for accelerated evolution of CA; the error bar is the posterior standard deviation. Colors as in (A). **C)** Accelerated evolution of CA for *DPP10* in humans. Colored, filled trace shows the moving average of the difference in the sum of human-derived and chimpanzee-derived predicted absolute log fold-changes. The black trace indicates one standard deviation above and below the mean absolute log fold-change difference of 100 iterations permuting which lineage each site was assigned to. **D)** Same as in (C) but in adult kidney proximal tubule cells. **E)** Genes with nominally accelerated evolution of CA (p < 0.05) in Purkinje cells. **F)** Same as in (A) but for sites with PhyloP > 1 and at least 250 species in the alignment. **G)** Enrichment of number of genes with nominal acceleration of CA (p < 0.05) in human vs. chimp. For each cell type, the bar on the left shows the enrichment value when restricting to only sites as described in (F) and the bar on the right shows the enrichment value when using all sites. Bars are red if there is an enrichment for the values for conserved sites relative to all sites (FDR < 0.05, Fisher’s exact test with Benjamini-Hochberg correction). **H)** Enrichments for human-derived substitutions in conserved sites in the top decile of accessibility (see Supp. Fig. 7F for cell type labels). **I)** Number of ASD-linked genes with nominally accelerated evolution of CA in at least one neuronal cell type. **J)** Same as in (I) but for sites with PhyloP > 1 and at least 250 species in the alignment. **K)** Schematic of a neuron with the subcellular localization of ASD-linked genes with nominal accelerated evolution of CA in conserved sites in at least one neuronal cell type.

To test whether lineage-specific selection on genes involved in ventricular septum morphogenesis may have impacted not only their 5’ UTR sequence evolution, but also their functional regulation, we examined their mRNA expression levels in cardiomyocytes derived from human-chimpanzee hybrid iPSCs^26^. These tetraploid hybrids, formed from *in vitro* fusion of human and chimpanzee iPSCs^27^, enable measurement of allele-specific expression which can only be caused by *cis*-regulatory divergence (including effects of UTRs). Strikingly, 13 out of the 14 tested genes had lower expression from the human allele (Fig. 2E-F, binomial FDR = 0.0036). This pattern is not consistent with equivalent selection pressures in both lineages (in which ∼50% of genes would be expected to have allelic bias in each direction), suggesting lineage-specific selection acting on the *cis*-regulation of these genes^28^. Notably, humans have considerable differences in heart morphology compared to other great apes^29^. As many of these genes are dosage sensitive^30^, their down-regulation may underlie some of these anatomical differences. Altogether, these results show that the excess of human acceleration extends to UTRs and is consistent with the idea that accelerated evolution can decrease gene expression.

### Accelerated evolution of the *cis*-regulatory landscape of genes and gene sets

We next turned our attention to substitutions in non-coding regions of the genome. As sequence alignment is more difficult in these regions, we used a set of strict filtering criteria to minimize artifacts due to poor alignment (see Methods). After filtering, we identified 25 human-accelerated genes (HAGs) and 11 chimpanzee-accelerated genes (CAGs, Fig. 3A-B, FDR < 0.05, Supp. Table 5). Again, even after controlling for the global difference in the number and conservation of substitutions (Supp. Text 3), there was an excess of human-accelerated genes (Fig. 3A, 2.3-fold more HAGs, p = 0.047, binomial test; this increases to 3-fold more HAGs with a more strict definition of acceleration, Supp. Text 1). Similar to the nonsynonymous and UTR analyses, most of the HAGs were driven by a shift toward substitutions in more conserved sites in the human lineage (12 out of 25 count-driven, 21 out of 25 weight-driven). However, the most significantly human-accelerated gene was entirely count-driven: *RBFOX1*, an RNA binding protein that plays a key role in neuronal development^31^, had a large excess of human-derived substitutions (Fig 3B-C) without any shift toward higher PhyloP scores (Supp. Fig. 5A).

**Figure 5:**
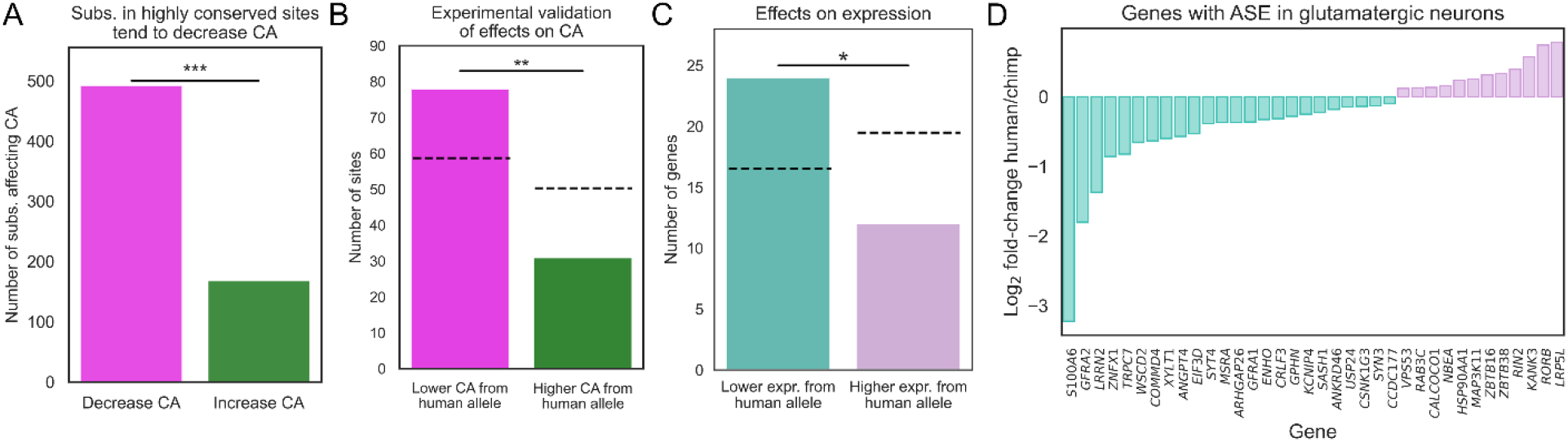
Human acceleration decreased chromatin accessibility. **A)** Substitutions in highly conserved sites (PhyloP > 6) that are predicted to decrease CA (predicted log_2_ fold-change < -0.5) or increase CA (> 0.5). *** p < 0.0005. **B)** Substitutions in conserved sites (PhyloP > 3) with predicted log_2_ fold-change < -0.5 that are in ATAC peaks with lower CA from the human allele (magenta) and higher CA from the human allele (green). Dashed lines indicate the number expected by chance. ** p < 0.005. **C)** Genes linked to an experimentally validated decrease in CA with lower expression from the human allele (teal) or higher expression from the human allele (purple). Dashed lines indicate the number expected by chance. * p < 0.05. **D)** Log_2_ fold-changes for genes from (C).

We then compared our results to previously identified HARs, as these are almost exclusively non-coding^4,6,7^. First, we intersected all our nominally significant (p < 0.05) HAGs and CAGs with a set of HARs aggregated across different studies^32^. If HAGs and HARs are capturing somewhat similar signals, we would expect HAGs to overlap with HARs more than CAGs. Consistent with this expectation, we found that the *cis*-regulatory neighborhoods of 21 nominal HAGs contained 3 or more HARs whereas this was true for only 6 CAGs (Supp. Fig. 5B, 3.3-fold enrichment, p = 0.0085, Fisher’s exact test). In contrast, there was no such enrichment for HAQERs, which are identified without regard to evolutionary conservation (1.2-fold depletion for genes with at least two HAQERs). Despite this overlap between HAGs and HARs, HAGs and HARs capture different signal and have different implications. First, 7 of our 25 genome-wide significant HAGs (*CXXC5*, *SIX3*, *NUMB*, *FTHL17*, *TMPRSS11F*, *FREM3*, and *MEOX2*) are not located near any known HARs^32^, suggesting that the accelerated evolution of their *cis*-regulatory landscape is entirely due to diffuse acceleration across conserved CREs near these genes. Second, even for HAGs that are linked to HARs, only a small fraction of substitutions in conserved sites are in HARs (Supp. Fig. 5C, 5.6% of human-derived substitutions in sites with PhyloP > 3 are in HARs for HAGs with at least one linked HAR). For example *RBFOX1*, despite having 17 nearby HARs (the fourth most of any gene^32^), this number is still only 5.6%. This suggests that accelerated evolution can extend broadly across many conserved CREs near specific genes and that many such CREs, even if they are not HARs, may have significantly accelerated functional divergence.

Searching HAGs for enriched functions, we found that transcriptional repressors were the only gene set with a significant bias toward human-acceleration relative to chimp (FDR = 0.015, Supp. Fig. 5D, Supp. Table 6). Testing genes categorized as both repressors and activators or exclusively repressors, there was significant acceleration only for those genes that only function as repressors (Fig 3C). Interestingly, many of these HAG repressors—including *SIX3*, *CXXC5*, *FOXP1*, *TCF7L2*, and *SCRT2*—play key roles in neurodevelopment^33–38^. For example, human-substitutions near *SIX3*, a transcriptional repressor essential for forebrain and eye development^38^, have higher PhyloP scores than those of chimpanzee-derived substitutions (Fig. 3E, Supp. Fig. 5E). To explore which individual *cis*-regulatory elements were driving the results for *SIX3*, we computed the FASTER test statistic in non-overlapping windows of 500 bases across the gene body and flanking sequence of *SIX3* (as well as all other human-accelerated genes; Supp. Table 7). The most accelerated region had 3 human-derived substitutions in conserved sites, including one that introduced a potential binding site for PAX6 (Fig. 3F), which regulates *SIX3* during eye development^39^. Collectively, these results suggest that transcriptional repressors experienced especially rapid evolution in conserved *cis*-regulatory elements during human evolution.

Given this connection to brain development and the complex expression patterns of many transcription factors during neurodevelopment, we next examined the expression of nominal HAGs and CAGs across different types of neurons. Using published snRNA-seq from the developing human neocortex^40^, we found that nominal HAGs were 3.1-fold enriched for being neuronal cell type-specific marker genes compared to genes that were not accelerated (p = 5.5x10^-6^, one-sided Fisher’s exact test for being among the top 25 marker genes per cell type) and enriched 2.0-fold relative to nominal CAGs (p = 0.048, Fig. 3G), suggesting that acceleration in the human lineage has a particularly strong impact on cell type-specific expression in the developing neocortex. These enrichments were largely driven by human-accelerated TFs that are essential for the differentiation of specific neuronal cell types such as layer 4 intratelencephalic neurons (L4 IT neurons) and are only expressed in one or a few cell types (Fig. 3H). For example, the nominal HAG *RORB*, a TF and marker for L4 IT neurons whose loss of function causes autism spectrum disorder (ASD)^41^, has decreased expression in humans relative to chimpanzees (FDR = 7.1x10^-6^) as well as other primates^42^ (Fig. 3I). Nominal HAGs are also 2.1-fold enriched for being genetically associated with ASD^41^ relative to unaccelerated genes (p = 0.0033, using non-syndromic genes from the SFARI database) and this enrichment persists when restricting to only loss of function intolerant genes (2.4-fold enrichment, p = 0.0028) suggesting that the signal for ASD is not the product of a general enrichment for loss of function intolerance. Moreover, HAGs are 2.2-fold enriched for ASD-linked genes relative to nominal CAGs (p = 0.037, Fig. 3J), primarily because nominal HAGs tend to be ASD-linked transcription factors (Fig. 3K) such as *RORB* and *FOXP1*^41^. Collectively, this suggests that specifically in humans, but not in chimpanzees, accelerated *cis*-regulatory divergence may have played a particularly important role in modifying the complex expression patterns of transcription factors that govern neocortical development as well as cognition more broadly.

### Human-biased accelerated evolution of chromatin accessibility only in conserved sites

A key aspect of FASTER is that it can test not only for accelerated evolution of sequence but also for accelerated evolution of function by using variant effect predictions to weight substitutions. In addition to conservation (used above), another important feature—which is central to transcriptional *cis*-regulation and possible to accurately predict from local sequence—is chromatin accessibility (CA). To scan for accelerated evolution of CA, we first generated predictions of the effects of human- and chimpanzee-derived substitutions on CA in 34 diverse cell types using the deep learning model ChromBPNet^43^. To enrich for active *cis*-regulatory elements, we restricted our analysis to the 10% of substitutions at positions with the highest predicted CA. Combining p-values from FASTER across cell types with mashr^44^ (while controlling for any genome-wide shift in predicted CA effect sizes; see Methods), we identified 142 genes with predicted human-accelerated CA evolution (CA-HAGs, 105 of which are weight-driven and 111 of which are count-driven) and 143 with predicted chimpanzee-accelerated CA evolution (CA-CAGs) (Fig. 4A, Supp. Table 8).

In general, signal was shared across many cell types. For example, *CSMD1*, which we have previously predicted to have uniquely rapid evolution of cell type-specific CA in the human lineage^45^, had human-accelerated CA in all but 3 cell types (Supp. Fig. 6). However, some genes had cell type-specific CA acceleration. For example, *DPP10*, a regulator of neuronal activity^46^, was significantly human-accelerated in Purkinje cells and pericytes but was chimpanzee-accelerated in committed oligodendrocyte precursors, adult kidney proximal tubule cells, and adult VIP+ cortical interneurons (Fig. 4B-D).

**Figure 6:**
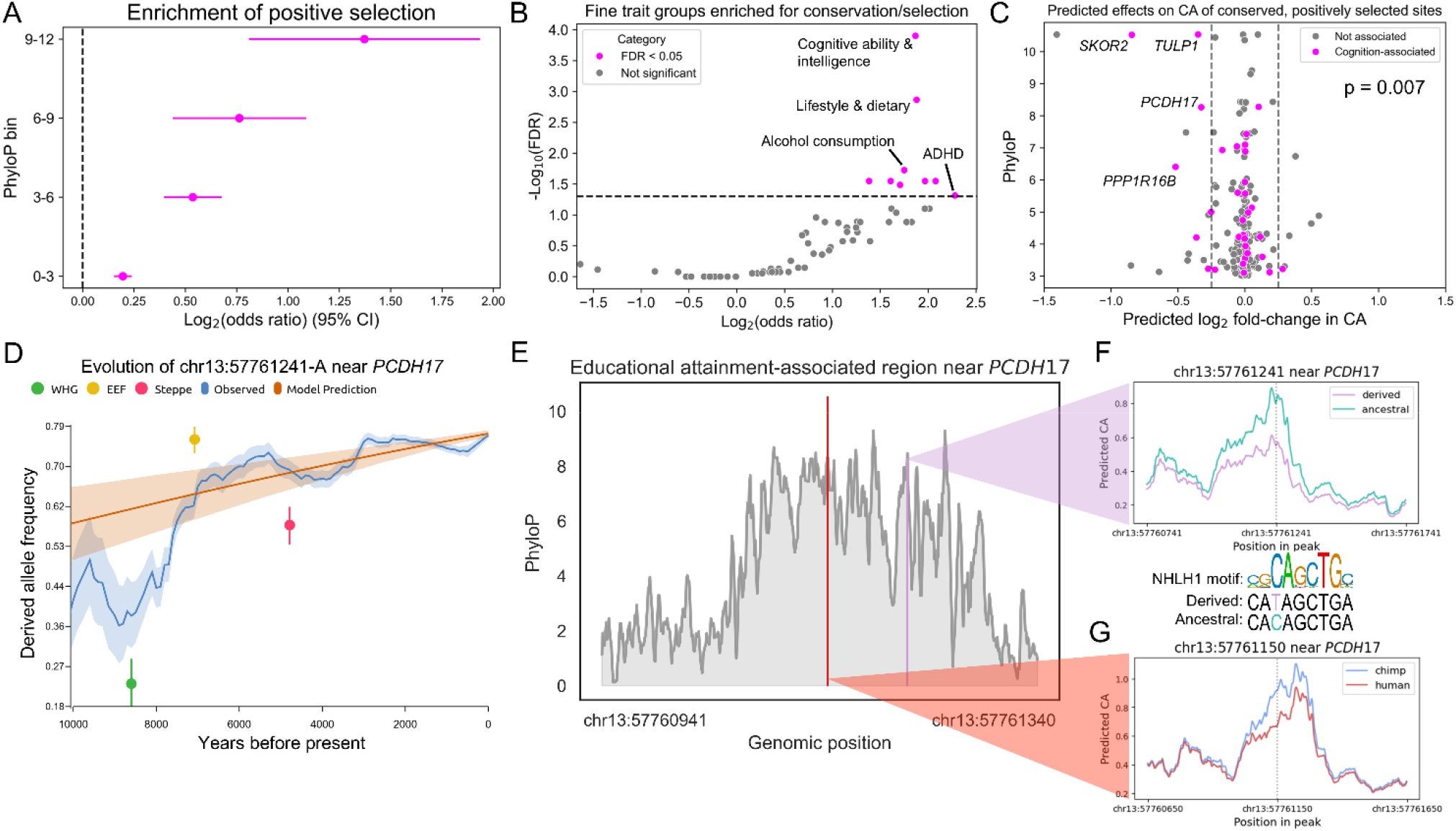
Disproportionate selection on conserved sites in modern human evolution. **A)** Enrichment of recently positively selected substitutions in conserved sites. **B)** Positively selected GWAS hit enrichments for highly conserved (PhyloP > 6) sites. Different GWAS were grouped by MeSH similarity. **C)** PhyloP scores (y-axis) and predicted effects on CA of the derived allele in fetal cortical neurons (x-axis) for sites with positive selection. Sites are colored magenta if they are associated with cognitive or psychiatric phenotypes. **D)** Derived allele frequency of chr13:57761241-A (rs1334297) over time in Europe. The blue trace is the observed frequency and the orange trace is the best linear fit under a constant selection coefficient. **E)** PhyloP scores for rs1334297 (purple) and a nearby fixed human-derived substitution (red). **F)** Predicted effect of the derived allele at rs1334297 on CA in fetal cortical neurons. NHLH1 motif (on the reverse strand), derived, and ancestral sequence are shown below. **G)** Same as in (F) but for the fixed substitution at chr13:57761150.

As mentioned above, we observed no difference in the number of CA-HAGs and CA-CAGs when either pooling information across cell types (Supp. Fig. 7A) or analyzing individual cell types (e.g. Purkinje cells in Fig. 4E). However, we reasoned that human acceleration could be specific to the most functionally important (and therefore conserved) sites. To test this hypothesis, we restricted to sites with PhyloP > 1 and repeated our analysis of accelerated evolution of CA. Remarkably, we observed an excess of genes with predicted human-accelerated evolution of CA in all 29 analyzed cell types (with 16/29 reaching statistical significance; Fig. 4F-G).

We noticed that many of the cell types with the strongest enrichments for human acceleration in conserved sites were neuronal (Fig. 4G), consistent with previous work showing that highly conserved non-coding elements tend to be active in the developing nervous system^47^. To explore this further, we calculated the enrichment of human-derived substitutions in highly conserved sites in predicted candidate CREs compared to non-CRE regions. Although all tested cell types showed some enrichment, the strongest enrichments were in the nervous system (19 out of the top 20 cell types were neurons, neural progenitors, or glia, with neurons showing the strongest enrichments; Fig 4H), suggesting that much of the accelerated CA evolution in conserved regions of the genome affects the nervous system.

To explore the potential consequences of this for ASD risk, we tested whether ASD-linked genes were more likely to have predicted human-accelerated or chimpanzee-accelerated CA evolution in three broad categories of cell types: neurons, glia/neural progenitors, and non-nervous system cells. Similar to the genome-wide results (Fig. 4E), we observed no overall enrichment for ASD-linked CA-HAGs in any cell type category (shown for neurons in Fig. 4I). However, conserved sites were enriched for human acceleration of CA for ASD-linked genes specifically in neurons (Fig. 4J, p = 0.015, binomial test), but not in other cell types (all p > 0.1, Supp. Fig. 7B-E). Interestingly, unlike ASD-linked HAGs (Fig. 3K), ASD-linked neuronal CA-HAGs were more likely to be involved in neuronal wiring and synaptic function than in transcriptional regulation (Fig. 4K), implying that two types of accelerated evolution—of the neuron-specific function of conserved sites vs. of the sequence of conserved noncoding sites more broadly—converge onto ASD-linked genes, despite affecting fundamentally different aspects of neurodevelopment. Overall, these results suggest that there is a bias toward more rapid functional evolution specifically in conserved regions of the human genome, and that this acceleration primarily affects specific genes that play important roles in neurodevelopment and cognition.

### Decreases in chromatin accessibility as a consequence of human accelerated evolution

Next, we wanted to investigate the consequences of human accelerated evolution on *cis*-regulation. As discussed above, one possibility is that this could generally result in decreased CA. To test this hypothesis, we stratified substitutions by their conservation level and compared the number of substitutions predicted to strongly increase or decrease CA. Consistent with our hypothesis, we observed a significant bias toward predicted decreased accessibility (Purkinje cells p = 9.0x10^-38^; Fig. 5A, see Supp. Table 9 for results across all 34 cell types) that became increasingly strong with greater conservation (Supp. Fig. 8A-D).

To experimentally validate these predictions, we examined published RNA-seq and ATAC-seq data from human-chimpanzee hybrid cortical neurons^48^. Using the ATAC-seq data (a direct measurement of CA), we confirmed that substitutions in conserved sites tend to decrease accessibility (Fig. 5B, 2.2-fold enrichment, p = 0.00086 for PhyloP > 3; Supp. Fig. 8E, 4.1-fold enrichment, p = 0.0087 for PhyloP > 6, Fisher’s exact test). Next, to explore the effects of these CA decreases on gene expression, we tested whether genes near the human-derived substitutions in conserved sites with large predicted negative effects on CA tended to have decreased expression from the human allele in hybrid neurons. We found a significant bias toward human down-regulation (Fig. 5C, p = 0.018, binomial test), including many genes that are essential for neuronal development. For example, both *GFRA1* and *GFRA2*, paralogous genes that interact with *RET* and play key roles in neuronal survival^49^, had decreased expression (Fig. 5D). Collectively, this supports the idea that one of the primary consequences of human *cis*-regulatory accelerated evolution is decreased CA, which often leads to decreased gene expression.

### Recent selection on conserved sites disproportionally affects cognitive traits

Next, we explored whether the trends we identified during hominid evolution extend to recent evolution in Europe using a published catalog of natural selection over the last 10,000 years inferred by estimating changes in allele frequency over time using ancient human DNA^50^. This enabled us to 1) examine if the human-acceleration we describe above has continued to impact recent human evolution, and 2) directly test whether human-acceleration was driven by positive selection, as opposed to other factors such as loss of constraint (see Supp. Text 1, Supp. Figs. 8-11 for additional analyses and discussion of alternative explanations for our results). Remarkably, we found that derived substitutions in conserved non-coding sites were enriched for positive selection and that this enrichment became progressively stronger with greater conservation (Fig. 6A, OR = 2.6, p = 2.2x10^-5^ for the most conserved sites; see also Supp. Fig. 12A). Derived alleles at high PhyloP sites were also enriched for negative selection, as is generally expected for conserved sites (Supp. Fig. 12B and Supp. Text 4). Collectively, these results suggest that substitutions in highly conserved sites have been subject to strong positive selection in the human lineage and this selection has continued to shape even very recent human evolution.

Unlike fixed substitutions, polymorphic substitutions can be directly linked to phenotype using genome-wide association studies (GWAS). To explore what traits are most affected by selection on substitutions in conserved sites, we intersected nominally selected (p < 10^-5^) sites with publicly available GWAS results^51^, grouping similar GWAS traits into broad categories to increase power and reduce redundant tests^52^. Using this strategy, positively selected highly conserved sites were enriched in several broad GWAS categories, particularly cognitive, psychiatric, diet, lifestyle, cardiovascular, and metabolic trait groups (Supp. Fig. 12C). In contrast, there were no enriched traits for negative selection (Supp. Fig. 12D). Using finer scale clustering of GWAS traits within the significant broad categories, the most significantly enriched trait groups were related to cognition (consisting of traits like “intelligence” and “educational attainment”), alcohol consumption, and several neuropsychiatric traits such as attention deficit hyperactivity disorder (ADHD) (Fig. 6B).

Given this enrichment for cognitive traits, together with the genome-wide trend for human-derived substitutions decreasing CA at conserved sites (Fig 5A-B), we hypothesized that recent positive selection at conserved sites may be biased toward decreasing CA specifically in fetal neurons. We found that positively selected derived alleles in conserved sites were indeed predicted to decrease CA in fetal cortical excitatory neurons more than expected by chance (Fig. 6C, OR = 2.64, p = 0.0011, one-sided Fisher’s exact test). Negatively selected variants were also somewhat enriched (Supp. Fig. 14E, OR = 2.19, p = 0.050, one-sided Fisher’s exact test). Notably, this trend was even stronger for cognition-associated variants (Fig. 6C, OR = 4.81 and p = 0.007, one-sided Fisher’s exact test) whereas neither derived alleles that increase CA nor derived alleles under negative selection that decrease CA were similarly enriched (all p > 0.6; Supp. Fig. 12E). Indeed, positively selected variants in conserved sites predicted to decrease CA were enriched for associations with cognition relative to negatively selected variants (7 positively selected associated with cognition, 0 negatively selected, p = 0.038). Combined with previous work showing that recent selective pressures tended to increase modern-day cognition^50^ (which we confirmed using our GWAS annotations, Supp. Fig. 12F-G), this suggests that the pattern of human-derived substitutions at conserved sites decreasing CA has extended into recent human evolution, where it was driven by positive selection that may have improved present-day human cognitive function (although there are important caveats for behavioral and cognitive GWAS, see Supp. Text 5). Overall, together with the lack of evidence of either reduced evolutionary constraint or potential confounding factors underlying the signals of acceleration we observe (Supp. Text 1), these results suggest that a considerable fraction of the predicted decreases in CA caused by human-derived substitutions may have been the result of positive selection (Table 1).

**Table 1:** Overview of models to explain our observations. The middle entry for “Reduced constraint in humans” is “Maybe” because while this would not be expected under a uniform, genome-wide decrease in constraint, it could happen if there were gene- or pathway-specific reductions in constraint. For detailed explanation of this table, see Supp. Text 1.

| **Observation** | No lineage-specific selection | Reduced constraint in humans | Positive selection in humans |
| --- | --- | --- | --- |
| Genome-wide bias toward human acceleration at conserved sites | No | Yes | Yes |
| Enrichment of acceleration in specific genes and pathways | No | Maybe | Yes |
| No excess of human polymorphism associated with acceleration | Yes | No | Yes |
| Recent positive selection enriched for substitutions in conserved sites | No | No | Yes |

As an example, we focused on a polymorphic substitution in a candidate CRE near *PCDH17*, which plays an important role in specifying neuronal connectivity^53,54^. The derived allele was under positive selection during recent human evolution, undergoing a dramatic increase in allele frequency around 7000-8000 years ago (Fig. 6D), despite this site being conserved in all but one of 446 other mammals. In addition, this substitution is predicted to decrease CA in fetal cortical neurons and disrupt the binding of NHLH1^55^, a transcription factor involved in specifying neuronal connectivity^56^ (Fig 6E-F). Most strikingly, the derived allele is one of the five strongest predictors of increased educational attainment genome-wide^57^ (p = 8x10^-90^). Despite this recent positive selection in Europe, the derived allele is at a comparable frequency across present-day human ancestry groups^58^ (Supp. Fig. 12H), suggesting either that this allele was recurrently positively selected in multiple ancestry groups or that ancestral Europeans experienced an initial decrease in allele frequency followed by selection to increase its frequency. Interestingly, this variant is in the same candidate CRE as a fixed human-derived substitution in a site that is conserved across all 446 other placental mammals (Fig. 6E). This fixed substitution is also predicted to decrease CA in fetal cortical neurons (Fig. 6G), suggesting that the recent positive selection occurred in the context of already reduced local CA that may also have altered human cognition.

## Discussion

Since the advent of comparative genomics, methods to detect accelerated evolution have had a major impact on our understanding of evolution across diverse lineages^4–7,59–68^. However, because of the inherent properties of these methods, they have mostly focused on the evolution of short, conserved (and more recently non-conserved^16^) non-coding sequences. Here, we developed a new framework to detect accelerated evolution that considerably expands on this strategy.

First, our method goes beyond sequence evolution by enabling the detection of accelerated evolution of any function encoded by DNA, RNA, or protein. For example, we identified over a hundred genes with human-accelerated evolution of chromatin accessibility, a key factor in *cis*-regulation. We expect that future applications of our method will enable researchers to detect and understand changes in the evolutionary rates of many different types of biological function, both in humans and across the tree of life. For instance, it is already possible to predict variant effects on properties including 3D genome conformation, transcriptional activity in reporter assays, protein stability, and UTR effects on gene expression^23,43,69–75^, and this list will undoubtedly continue to grow.

Second, our method can be applied to any set of genomic regions, including (but not limited to) proteins, UTRs, and *cis*-regulatory neighborhoods of genes and pathways. This flexibility enables the investigation of accelerated evolution at many different levels of genomic organization. Applying our method to humans and chimpanzees, we identified dozens of human-accelerated proteins (HAPs), UTRs (HAUs), and *cis*-regulatory neighborhoods of entire genes (HAGs). These included three neurodevelopmental HAPs (RET, ESR1, and EYA1) with human-derived changes in their stability, degradation, or processing—changes that could impact protein activity levels, not unlike *cis*-regulatory changes to transcription or translation, but through a distinct mechanism. In the future, it will be interesting to explore the phenotypic impacts of the substitutions in HAPs, HAUs, and HAGs. We anticipate that such efforts will help uncover the genetic basis of uniquely human traits, in much the same way as they have for HARs^32,76–78^.

Across diverse analyses, we consistently found that accelerated evolution of sequence and chromatin accessibility in conserved non-coding sequences – both throughout the ∼6 million years of hominid evolution as well as in modern humans – disproportionately occurred in the human lineage, were predicted to reduce function, and primarily affected genes involved in neurodevelopment, neuropsychiatric disorders, and general cognition. Notably, we found that substitutions in conserved sites under recent positive selection – especially those predicted to decrease CA in developing neurons – were enriched for cognitive and psychiatric traits in GWAS. This, combined with other analyses that failed to find evidence for loss of constraint, suggests that a considerable proportion of the signal of accelerated evolution we observe was due to positive selection (although relaxed constraint could still play a role in driving the genome-wide pattern of human acceleration). This provides a potential molecular mechanism to help explain our recent finding that positive selection for decreased gene expression of ASD-linked genes played an important role in the evolution of human brain development and cognition^14^. Collectively, our results paint a coherent picture of human evolution in which positive selection for reduced function accelerated evolutionary innovation across diverse conserved regions of the genome. Overall, this suggests that adaptive decreases in *cis*-regulatory activity—and the resulting down-regulation of a wide swath of mRNAs and proteins—was an important component of the evolution of uniquely human cognition, with broad implications for understanding the genetic basis of uniquely human traits and disease susceptibility.

## Supporting information

Supplemental Zip File 1

## Glossary

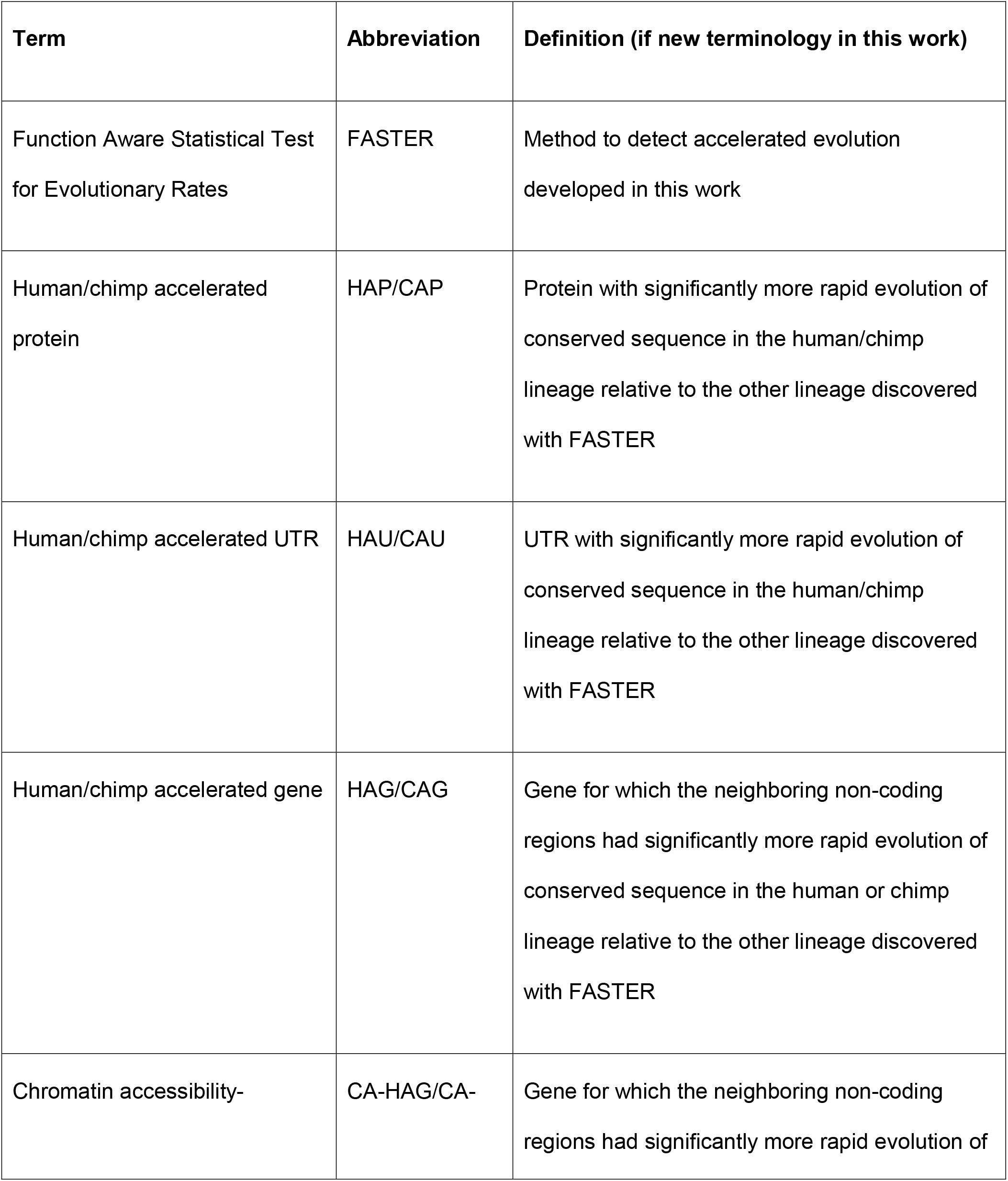
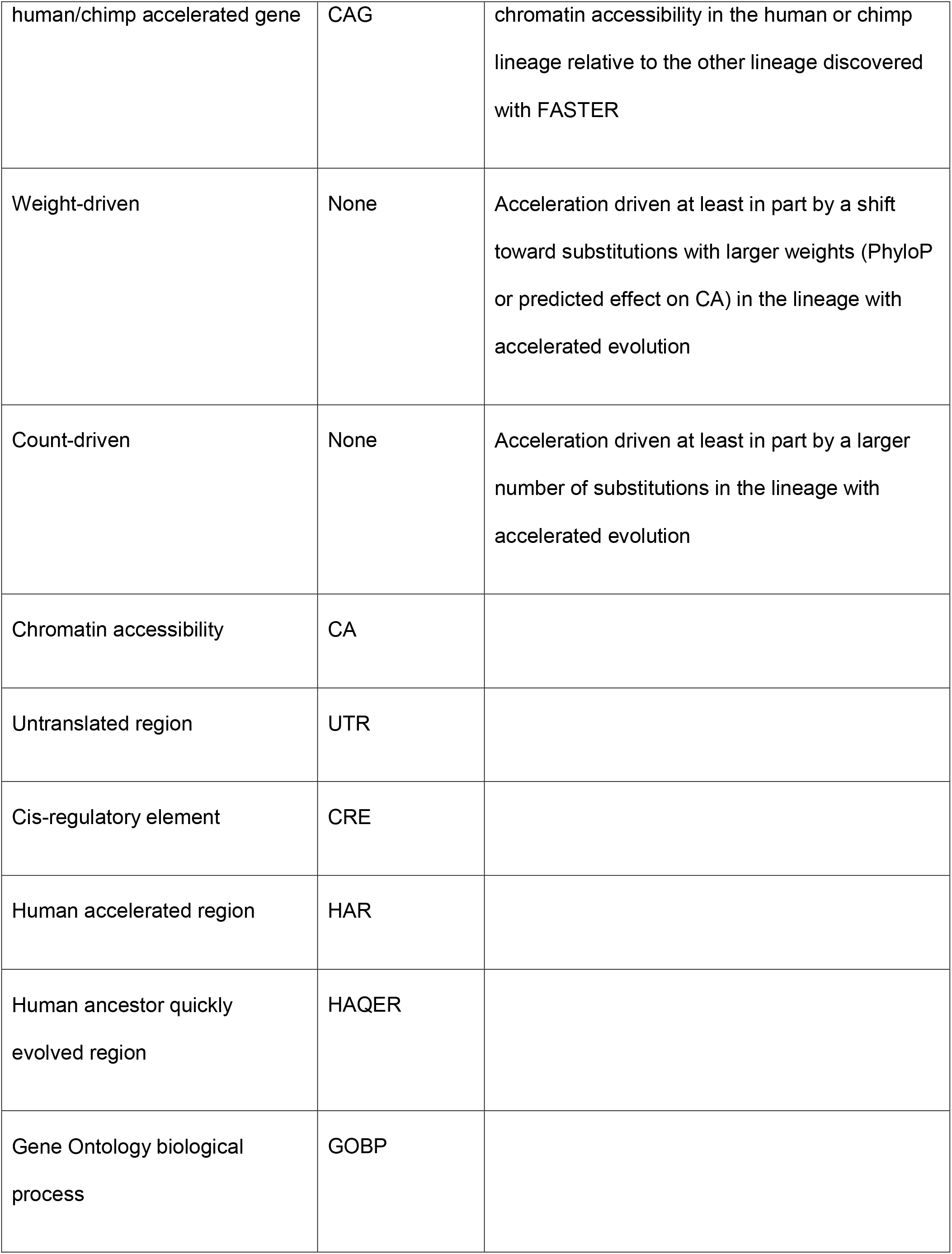
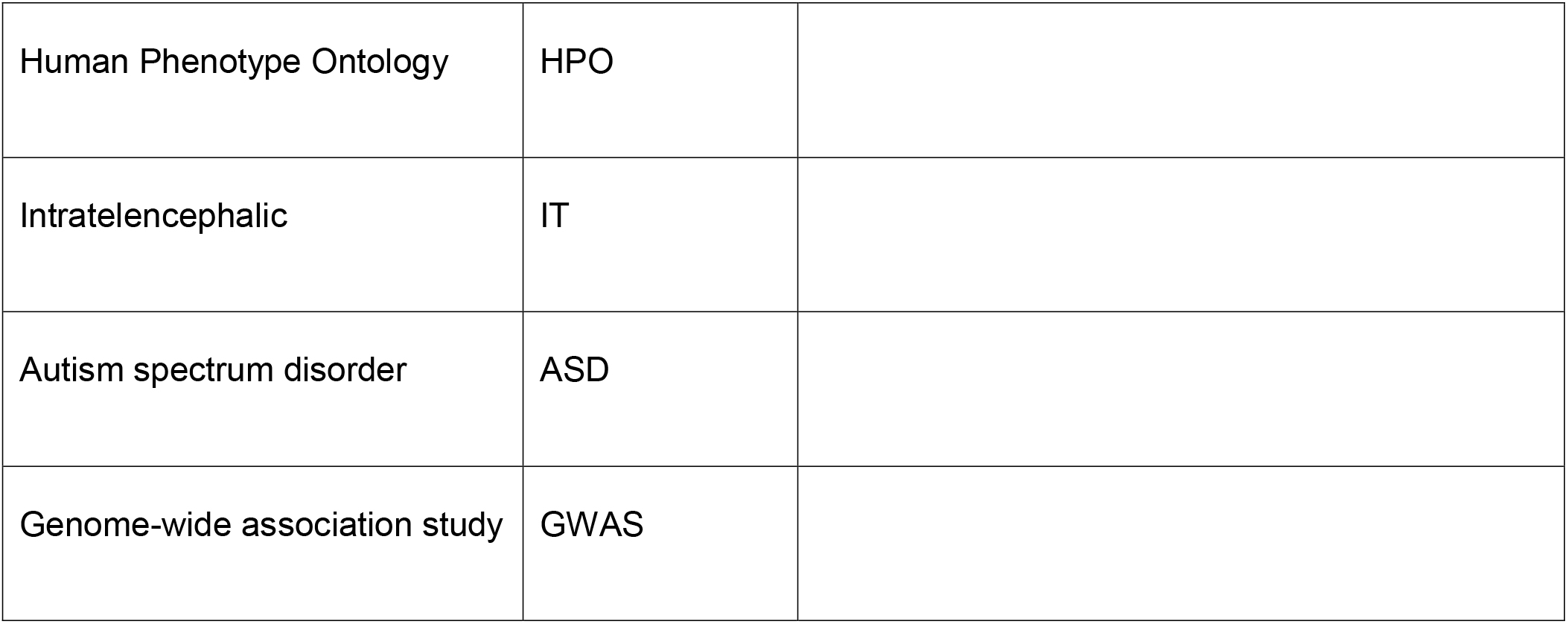

## Methods

### Generation of the input dataset for testing for accelerated evolution

To identify and classify substitutions, we used our previously published approach^26,27^. Briefly, we first used the Hg38-PanTro6 whole genome alignment to identify substitutions and compared to the GorGor6 gorilla genome to determine if a substitution was human- or chimpanzee-derived. If the gorilla base did not match the human or chimpanzee base, we discarded it. We then assigned substitutions as nonsynonymous, 3’ UTR, 5’ UTR, or noncoding using ENSEMBL variant effect predictor v109^79^. All substitutions altering amino acid sequences were classified as nonsynonymous, all in the 3’ UTR as 3’ UTR unless they were also classified as nonsynonymous, all in the 5’ UTR as 5’ UTR if they were not classified as nonsynonymous or 3’ UTR, and as non-coding if they were not classified as any of the other categories.

We used our previously published approach^26^ to filter out polymorphisms in both lineages and create a high-confidence set of substitutions by checking whether 3 human iPSC and 3 chimpanzee iPSC lines were homozygous for their respective reference alleles. We use this filtered set for the nonsynonymous and UTR analyses and both the filtered and unfiltered versions separately for the non-coding analyses (see below). For all analyses, we only included genes that were three-way one-to-one orthologs across human, chimpanzee, and gorilla defined using ENSEMBL biomart v109^80^. We also created a blacklist of genes near centromeres and/or with cross-species alignment problems.

We used our previously published approach in conjunction with phast v1.5^5,11,81^ to compute PhyloP scores while masking the human, chimpanzee, and bonobo genomes. When using PhyloP to test for accelerated evolution, we replace all negative values with zero to avoid penalizing substitutions in rapidly evolving sites. We additionally computed the number of genomes in the alignment at each position and filtered to only substitutions in sites with at least 250 species in the alignment for all PhyloP-based analyses to ensure that there was sufficient statistical power to estimate conservation as previously described^45^. We used our previously published set of ChromBPNet models for each of 34 cell types (see Starr et al. 2025^45^ for the complete list of cell types and abbreviations) to predict the effects of reverting human-derived substitutions to the ancestral base in the human sequence context and reverting chimpanzee-substitutions to the ancestral base in the chimpanzee sequence context to ensure the two sets of variant effect predictions were directly comparable^45^. We assigned non-coding substitutions to the nearest protein-coding transcription start site and considered the genomic region between the leftmost and rightmost substitutions assigned to a gene to be the *cis*-regulatory neighborhood of that gene^45^.

### General methodology to test for accelerated evolution with FASTER

In line with the core methodology of previously published methods to detect accelerated evolution^4,6,7^, we started by computing the weighted sum of substitutions in one lineage (in this case human) compared to a sister lineage (chimpanzee). We then calculated the difference between these two values as our test statistic. It is worth noting that many different test statistics and weighting schemes are possible. For example, one could estimate each variant’s selection coefficient as its weight. In practice, we selected the difference-of-sums test statistic as it was not overly affected by a few outlier substitutions with very high PhyloP scores or large effects on CA and because it is, in principle, similar to previous test statistics used to detect accelerated evolution. To assign a p-value, we implemented a sampling procedure in which each observed substitution was first randomly assigned to one of the two lineages and then to a randomly selected gene (or gene set). For the former, this assignment was performed according to a binomial distribution parametrized by the observed genome-wide probability that a substitution in that particular class occurred in the human or chimpanzee lineage. For the latter, we used a multinomial distribution over all input genes (or gene sets). Here, the class of a substitution is flexibly defined by some set of properties depending on the particular analysis being performed and what factors we want to control for. For example, to control for potential differences in the mutational profile between species in non-coding analyses, we defined the class of each substitution by the ancestral trinucleotide context of a site, in effect using a separate probability distribution for each of the 64 possible ancestral trinucleotide contexts. Continuing this example, all substitutions that occurred at the G in an ancestral AGC were assigned one probability, those that occurred at the central A in an ancestral AAC a different probability, etc. Below, we detail how this class was defined for each analysis and the exact details of how p-values were computed. When testing directly for accelerated evolution of gene sets, we used an identical procedure to assign to genes and then compute the actual or sampled test statistic for the gene set. When testing for accelerated evolution of genes in GOBP or HPO gene sets, we restricted to sets with between 20 and 200 genes. We considered a gene to be count-driven if there were nominally significantly (i.e. p < 0.05) more substitutions in the accelerated lineage by a binomial test and weight-driven when the median of the distribution of PhyloP scores was significantly greater in the accelerated lineage (Mann-Whitney U test p < 0.05). The exception to this was for non-coding regions, for which we used an extension of FASTER to test whether a gene was weight-driven.

### Testing for accelerated evolution of protein-coding regions

For protein-coding regions, we defined the class as the ancestral amino acid to control for biases in nonsynonymous substitution rate that may not be reflected in DNA sequence alone (this is what prevents us from controlling for the global shift toward higher PhyloP scores in the human lineage for protein-coding analyses; PhyloP is a per-nucleotide statistic whereas our class is defined at the level of amino acids). We restricted to genes with at least 3 observed nonsynonymous substitutions in either the human or chimpanzee lineage. We then performed our sampling procedure one million times and computed a p-value for each gene as the number of samples with a test statistic more extreme than the observed test statistic divided by the total number of samples multiplied by 2. We computed an FDR and corrected for multiple testing using the Benjamini-Hochberg method. For all analyses, we tested for more acceleration in the human or chimpanzee lineage using the binomial test. We repeated our per-gene analysis for Gene Ontology biological process terms (GOBP) and Human Phenotype Ontology terms (HPO) using all genes as input. To identify enrichments that were not driven by outlier proteins (e.g. RET), we removed the top three most human- and chimpanzee-accelerated genes from each term. We used an independent t-test to test for differences in the distribution of PhyloP scores for human- and chimpanzee-derived substitutions in GOBP and HPO terms.

To investigate the effects of substitutions in conserved sites on protein stability, we intersected our set of human-derived nonsynonymous substitutions with the predicted effects on protein stability from RaSP (which does not use conservation as an input for its predictions)^23^. We considered a substitution as destabilizing if it had predicted ΔΔG > 2 and non-destabilizing if it had ΔΔG < 0. We then binned sites by PhyloP score and used Fisher’s exact test to determine whether the substitutions in that bin were enriched for destabilizing substitutions relative to those with PhyloP < 0.

### Testing for accelerated evolution of UTRs and further analysis

To test for accelerated UTR evolution, we separated 5’ and 3’ UTRs. We defined the class of a substitution according to the ancestral trinucleotide context in which the substitution occurred and, only for assigning substitutions to each lineage, the PhyloP score of the site. For the latter, we assigned substitutions to one of one hundred bins of equal width, ranging from the minimum to the maximum PhyloP score. We then computed a p-value and FDR for genes and gene sets in the same manner as for nonsynonymous substitutions. To test for lineage-specific selection on genes involved in ventricular septum morphogenesis, we used previously published allele-specific expression (ASE) data from human-chimpanzee hybrid iPSC-derived cardiomyocytes (see Wang et al. for details on processing of this data, which included correction for mapping bias^26^). We used a binomial test parameterized by the background probability of a gene having lower expression from the human allele across all genes with ASE (at FDR < 0.05, as we tested all GOBP categories with significant 5’ UTR acceleration) to test whether there were significantly more ventricular septum morphogenesis genes with lower expression from the human allele than expected by chance.

### Testing for accelerated *cis*-regulatory evolution

For *cis*-regulatory neighborhoods, we used a similar strategy as for UTRs, defining the class of each substitution in the same way. However, as there are many more non-coding substitutions, we performed 1000 samplings to produce a null distribution and then computed a z-score and p-value using the normal approximation. We also ran the test twice, once with our filtered set of substitutions and once with the unfiltered set as we noticed that for large non-coding regions near several genes, the filtering process removed an unexpectedly large number of chimpanzee-derived substitutions (see below for how we used p-values from both tests in identifying HAGs). In addition to the p-value computed in the same manner as for UTRs, we also computed two additional p-values. First, to correct for potential biases in how substitutions were assigned to genes, we scaled the summed human-derived and chimpanzee-derived substitutions for each sampling to what it would have been had our sampling assigned that gene the same total number of substitutions as we observed. This was necessary because a small group of genes had very different numbers of substitutions assigned during sampling than occurred in reality.

To see this, consider an extreme example in which a gene with 1000 observed substitutions had only 100 assigned to that gene on average during sampling. If the sum of non-negative PhyloP were 550 for human and 500 for chimp, then our test statistic would be 50. Even if this test statistic were expected based on the number of substitutions that occurred genome-wide in the human and chimpanzee lineages respectively, the average sampling would produce a PhyloP sum of 55 for human, 50 for chimp, and a test statistic of 5 (as there are 1/10^th^ as many substitutions in the average sampling). This could lead to an artificially anti-conservative z-score. To correct for this, for each lineage in each sampling, we multiplied by a correction factor: the total number of observed substitutions divided by twice the number of sampled substitutions for the lineage. Continuing our example, if 50 substitutions were assigned to the human lineage for a particular sampling, our correction factor would be 1000/(2*50) = 10. Multiplying our sampled test statistic of 55 by 10 leads to 550, which is expected if the observed value for the human lineage of 550 was due entirely to genome-wide differences between human and chimp. The converse scenario could also be true, in which not performing this correction led to an artificially conservative z-score. In practice, we found that this latter scenario is likely more prevalent. However, to be conservative, we required an FDR < 0.05 for both the uncorrected and total corrected versions of the test. Without incorporating the total corrected p-value, we observed a similar but slightly more significant bias towards human-acceleration (29 HAGs vs. 13 CAGs; p = 0.019, binomial test).

To ensure that any genes we identified were not driven by technical factors, we filtered to only genes with at least 100 human-derived or chimpanzee-derived non-coding substitutions and considered a gene to be significantly accelerated if it had an unfiltered FDR < 0.05, an unfiltered total corrected FDR < 0.05, a filtered FDR < 0.25, and a filtered total corrected FDR < 0.25. This filtering excluded genes that were called as accelerated as a result of mismatches between our sampling and observations (as described in the preceding paragraph), those with acceleration primarily driven by substitutions in difficult to align or polymorphic sites (as a result of the substitution filtering step based on human and chimpanzee whole genome-sequencing), and those for which unexpected technically-driven consequences of our whole genome-sequencing based filtering drove the signal for acceleration, leaving us with a conservative set of accelerated genes.

Rather than use the Mann-Whitney U test to determine what acceleration was weight-driven (which would give undue weight to the many non-coding substitutions with low, positive PhyloP scores), we used an extension of FASTER to test for shifts in the distribution of PhyloP scores. We did this by normalizing the human-derived and chimpanzee-derived sums by their respective sampled number of substitutions divided by the actual number of substitutions observed for that gene in each lineage (referred to from here on as the distribution p-value). For example, if a gene had 600 observed human- derived substitutions and 400 observed chimpanzee-derived substitutions and, during a sampling, 500 substitutions were assigned to the human lineage and 500 assigned to the chimpanzee lineage, we would multiply the human sum of non-negative PhyloP by 500/600 and the chimpanzee sum by 500/400. As a result, the total number of substitutions assigned to the human lineage in each sampling would always be 1.5x greater than the number assigned to the chimpanzee lineage, effectively removing the effect of differences in the number of substitutions and only testing for a shift in the distribution of PhyloP scores. We used this test only to determine the extent to which results from the preceding paragraph were weight-driven, not to classify genes as HAGs or CAGs (so genes like *RBFOX1* are classified as count-driven HAGs). We used a binomial test to determine which genes were count-driven.

For the analysis of repressors and activators, we considered a gene to be a repressor if it was in the gene set associated with the GOBP term Negative regulation of DNA-templated transcription and not in the term Positive regulation of DNA-templated transcription (and *vice versa* for activators). We then used an independent t-test against all genes not in either GOBP term to test for a shift in the distribution of PhyloP scores. For testing across all GOBP terms, we used each term with at least 20 genes assigned in conjunction with an independent t-test.

To define nominal HAGs and CAGs, we required p < 0.05 and total corrected p < 0.05 for both the filtered and unfiltered set of substitutions. For the analyses of enrichment for ASD-linked genes, we used all SFARI^41^ genes and Fisher’s exact test, restricting to nominal HAGs, nominal CAGs, or genes that were not nominal HAGs or nominal CAGs. We used this same strategy for the genes with nominal acceleration of chromatin accessibility. ASD-linked genes were annotated as having synaptic or nuclear functions using SYNGO^82^, GO^21^, and manual curation. For the developing cortical markers, we used data from Wang et al.^40^. We normalized the raw snRNA-seq, restricted to only neuronal cell types defined by Wang et al., computed the mean expression across cells within each neuronal type, computed the fold-change of expression in each cell type over the mean expression across all other neuronal types, ranked genes by this metric, and then took the top 25 genes for each neuronal type as marker genes for that neuronal type. We then used Fisher’s exact test to test for enrichment in each neuronal type separately and for all genes that were a marker gene in at least one neuronal type.

For CA with no filtering by site conservation, we used this same strategy with the following exceptions. First, we used any site, regardless of the number of species in the alignment for that site. Second, for each cell type separately, we restricted to only substitutions in the top decile of total CA, where total CA was the maximum CA of the human and chimpanzee allele for each site. When first filtering by site conservation, we used the same strategy except that we required that an input site have at least 250 species in the alignment and PhyloP > 1. For mashr^44^, we input the acceleration z-scores for filtered/unfiltered and total corrected/non-total corrected p-values. We then restricted to only genes that had a local false sign rate (LFSR) from mashr less than 0.05 for both filtered and unfiltered or LFSR greater than 0.05 for both filtered and unfiltered and then reran mashr on the remaining genes. We then applied the same filtering criteria as the PhyloP analysis described above, replacing FDR with LFSR.

### Visualization of accelerated evolution

To visualize accelerated evolution in single CDSs, UTRs, and 500 base non-coding windows, we plotted the moving average of PhyloP for each site in that region. We then overlaid red lines for human-derived substitutions and blue lines for chimpanzee-derived substitutions. To visualize the *cis*-regulatory neighborhood of genes, we computed a moving average of our test statistic computed in 500 base pair windows tiling the *cis*-regulatory neighborhood of a gene. We then permuted which lineage (human or chimp) each substitution was assigned to, recomputed the moving average 100 times, and then plotted the standard deviation of the result. For CA, we used an identical procedure, except that we stitched together all substitutions in the top decile of accessibility (rather than fully tiling the region). For plots of PhyloP distributions, we used seaborn^83^ v0.13.2 kdeplot, normalized such that the area under the curve was identical for all distributions. The genomic positions for introns and exons were for the ENSEMBL v109 canonical transcript.

### Analysis of ILS and GC-biased gene conversion (gBGC)

To identify substitutions consistent with ILS, we scanned the genome for sites that support alternative tree topologies (either chimpanzee more closely related to gorilla [CGH] or human more closely related to gorilla [HGC]; the former were considered human-derived in the main analysis and the latter chimpanzee-derived). We define CGH sites as those where chimpanzee and gorilla have the same base, human has a different base, yet orangutan has the same base as human (and *vice versa* for HGC sites)^84^. We then computed the weighted non-negative sum of PhyloP scores for nonsynonymous, 3’ UTR, 5’ UTR, and non-coding substitutions and the difference between the human and chimpanzee lineages as described above. We computed the fraction of acceleration explainable by ILS as the FASTER test statistic (i.e. the difference in weighted sum between the human and chimpanzee lineages) for ILS sites alone divided by the FASTER test statistic for all sites. Negative values were clipped to zero and values greater than 1 were clipped to 1 (these situations occurred when removing ILS sites increased the signal of acceleration). In addition, we removed all ILS sites and reran the test for a genome-wide bias toward substitutions in higher PhyloP sites in the human lineage. We used the same strategy to analyze gBGC, using a previously published catalog of gBGC regions in human and chimpanzee^85^. We lifted (using UCSC liftover) the gBGC regions from hg18 to hg38 and considered any human-derived A/T to G/C substitution in a human gBGC region or chimp-derived A/T to G/C substitution in a chimp gBGC region to be caused by gBGC.

### Testing for biases in ChromBPNet predictions

We computed the mean absolute predicted log fold-change for human-derived and chimp-derived substitutions separately for all sites and those with PhyloP > 1. We computed the bias toward substitutions decreasing accessibility in more conserved sites for chimpanzee-derived sites in the same manner as human-derived sites.

### Analysis of the potential role of lineage-specific deceleration

Throughout our primary analyses, we refer to a greater PhyloP-weighted substitution signal in one lineage relative to another as lineage-specific acceleration. However, a difference between two lineages could theoretically arise either through increased substitution at conserved sites in one lineage (acceleration), or decreased substitution at conserved sites in the other lineage (deceleration). To distinguish between these possibilities, we used the gorilla lineage as an outgroup and compared the PhyloP-weighted substitution signal across the human, chimpanzee, and gorilla lineages.

We identified gorilla-derived substitutions as those sites for which the gorilla base differed from the human, chimpanzee, and orangutan bases and all three bases from the non-gorilla species were the same (using the Hg38-PonAbe3 multi-way alignment and the same methodology outlined above for human-derived and chimp-derived substitutions). For the protein-coding region, 3’ UTR, 5’ UTR, and non-coding *cis*-regulatory regions of each gene, we calculated their PhyloP-weighted sum using the same genomic regions and variant classes used for the corresponding human–chimpanzee analysis. Because the gorilla lineage has a longer branch length than either the human or chimpanzee lineage and therefore contains more lineage-specific substitutions, we normalized the gorilla PhyloP-weighted sums before making gene-level comparisons. We first calculated the genome-wide total PhyloP-weighted sum for each lineage. The expected human/chimpanzee lineage total was defined as the mean of the genome-wide human and chimpanzee totals. We then calculated a genome-wide gorilla scaling factor as the ratio of the total gorilla PhyloP-weighted sum to this human/chimpanzee mean. Each gene’s gorilla PhyloP-weighted sum was divided by this factor, placing the gorilla signal on the same genome-wide scale as the human and chimpanzee signals.

We then used the normalized gorilla value to decompose each human–chimpanzee difference into components attributable to acceleration of the lineage with the greater PhyloP-weighted signal and deceleration of the lineage with the lower signal. For genes with a greater human than chimpanzee PhyloP-weighted sum, the fraction attributable to human acceleration was calculated as the difference between the human and normalized gorilla sums divided by the human–chimpanzee difference. The fraction attributable to chimpanzee deceleration was calculated as the difference between the normalized gorilla and chimpanzee sums divided by the human–chimpanzee difference. Conversely, for genes with greater chimpanzee than human signal, the acceleration fraction was calculated from the chimpanzee–gorilla difference and the deceleration fraction from the gorilla–human difference, each divided by the chimpanzee–human difference.

When the normalized gorilla value fell between the human and chimpanzee values, these two components sum to one and provide a direct decomposition of the observed human–chimpanzee difference. A normalized gorilla value outside the interval defined by the human and chimpanzee values can produce a negative fraction for one component and a value greater than one for the other, indicating that the outgroup comparison is not consistent with a simple mixture of acceleration in one lineage and deceleration in the other. For summary and visualization purposes, acceleration and deceleration fractions were therefore independently clipped to the interval from zero to one. Thus, a clipped acceleration fraction of one indicates that the outgroup comparison is consistent with the observed difference being entirely attributable to acceleration of the faster-evolving lineage, whereas a value of zero indicates that it is consistent with deceleration of the other lineage. Intermediate values indicate contributions from both processes. Throughout the main text, we report the ratio of human- to chimpanzee-accelerated genes, proteins, or UTRs obtained when restricting to only genes, proteins, or UTRs with an acceleration fraction greater than 0.9 in addition to that obtained with no filtering based on acceleration fraction.

### Selection, reconstruction, and cloning of human and LCA proteins

To select HAPs for experimental testing, we focused on proteins that were at least nominally human-accelerated, less than 2100 amino acids in length (to facilitate cloning), were haploinsufficient (i.e. gnomad probability of loss of function intolerance > 0.9; to enrich for proteins likely to affect phenotypes), and, ideally, had good commercially available antibodies (although this was not a hard requirement). Therefore our results may not be representative of HAPs which are longer in length or not haploinsufficient. To reconstruct the last common ancestor (LCA) allele of each protein, we reverted each human-derived amino acid change to the amino acid shared at that position by gorilla and chimpanzee. The human and reconstructed LCA genes were then codon optimized, ordered as gene fragments from Twist Biosciences (South San Francisco, CA), and cloned together in our plasmid backbone using the NEBuilder® HiFi DNA Assembly Cloning Kit (cat no. E2621S, New England Biolabs, Ipswich, MA). EYA1 was N-terminally FLAG tagged^86^ and TSHR and AKAP11 were C-terminally FLAG tagged^87,88^. All plasmids except GFRA1 also express mCherry that is co-translated with, but not translationally fused to, the gene of interest. Complete plasmid information can be found in Supplemental Zip File 1.

### Cell lines and culture conditions

MDA-MB-231 (cat no. HTB-26) and 293T (cat no. CRL-3216) cell lines were obtained from American Type Culture Collection (Manassas, VA). MDA-MB-231 cells were grown in Dulbecco’s Modified Eagle Medium (DMEM, cat no.10569010, Thermo Fisher, Waltham, MA), 10% charcoal-stripped fetal bovine serum (cat no. A3382101, Thermo Fisher), and 0.5 U/mL penicillin-streptomycin (P/S, cat no. 15070-063, Thermo Fisher). 293T cells were grown in DMEM, 10% FBS (cat no. 16000044, Thermo Fisher), and 0.5 U/mL P/S. Trypsin (cat no. 5200072) was from Thermo Fisher. Phosphate buffered saline (cat no. P5368) was from Millipore Sigma (St. Louis, MO). For all experiments, cells were trypsinized and counted on a Countess II Automated Cell Counter (cat no. AMQAX1000, Thermo Fisher). For all experiments, cells were grown and treated in a humidified 37°C, 5% CO_2_ incubator unless otherwise indicated. 293T cells were used for most proteins due to technical feasibility and the established role of EYA1, AKAP11, and RET in kidney function and development^89–91^. MDA-MB-231 cells were used as they are an ESR1-negative breast cancer model. It is unknown whether the effects on steady-state protein abundance observed here would also be observed in other cellular contexts.

### Western blotting

The day before the experiment: for RET, 100,000 293T cells/well were seeded into a 12-well (cat no. 07-200-82, Thermo Fisher) plate; for EYA1, TSHR, and AKAP11, 300,000 293T cells/well were seeded into a 6-well plate (cat no. 08-772-1B, Thermo Fisher); for ESR1, 350,000 MDA-MB-231 cells/well were seeded into a 6-well plate (Thermo Fisher). All cells were seeded in appropriate antibiotic-free growth medium.

The following day, cells were transfected using TransIT-LT1 Transfection Reagent (cat no. 2304, Mirus Bio, Madison, WI) according to the manufacturer’s instructions. 293T cells were co-transfected with GFRA1 (0.25 µg) and either HUM_RET (0.25 µg) or LCA_RET (0.25 µg) per well. All other plasmids were transfected at 2 µg/well. Cells were incubated for 48 h (MDA-MB-231) or 72 h (293T). After the incubation, cells were washed once with ice-cold 1x PBS and harvested in 0.5-1 mL of ice-cold 1x PBS by scraping. Cell suspensions were transferred to sterile microcentrifuge tubes and pelleted by centrifugation (300 × g, 5 min, 4°C). Cell pellets were washed once with 1 mL ice-cold 1x PBS and re-pelleted by centrifugation (300 × g, 5 min, 4°C). After removal of supernatant, pellets were either processed immediately or stored at -80°C until needed.

On the day of cell lysis, cell pellets were resuspended in RIPA buffer (cat no. PI89900, Thermo Fisher) supplemented with protease inhibitor cocktail (cat no. PPC2020, Sigma-Aldrich), 1 mM PMSF (cat no. PI36978, Thermo Fisher), and 5 mM NaF (cat no. S6776, Millipore Sigma). 60-100 µL of lysis buffer was used per well of a 6-well plate, depending on pellet size. After resuspension, cell suspensions were incubated on ice for 45–60 minutes, followed by sonication [(1 s on, 1 s off, 20-40% amplitude) x 10 cycles]. Lysates were clarified by centrifugation (13,800 × g, 15 minutes, 4°C). Lysates were quantified using a Pierce BCA assay kit (cat no. 23225, Thermo Scientific) with a standard BSA curve, according to the manufacturer’s instructions. Absorbance (562 nm) was measured using a Cytation 3 multimode plate reader (BioTek, Winooski, VT).

Equal amounts of protein were combined with 10x Bolt Sample Reducing Agent (cat no. B0009, Thermo Fisher) and 4x Bolt LDS Sample Buffer (cat no. B0007, Thermo Fisher) and then loaded onto a Bolt 4–12% Bis-Tris Plus Gel (cat no. NW04120BOX, Thermo Fisher). A lane of Chameleon Duo pre-stained protein ladder (cat no. 92860000, LI-COR Biosciences, Lincoln, NB) was run on every gel to facilitate gel migration, orientation, and estimation of protein size. Lysates from MDA-MB-231 cells were run in 1x MES buffer (cat no. B0002, Thermo Fisher). Lysates from 293T cells were run in 1x MOPS buffer (cat no. B0001, Thermo Fisher). Protein was transferred to a nitrocellulose membrane using an iBlot2 transfer stack (cat no. IB23001, Thermo Fisher). The membrane was blocked using Intercept Blocking Buffer (cat no. 927-60001, LI-COR Biotechnology, Lincoln, NE) (60 rpm, 1 h, room temperature) and then incubated in the appropriate primary antibody mixture (4°C, overnight). All primary antibodies were diluted in Intercept (LI-COR). Primary antibodies used were: α-ESR1 (cat no. 8644S, Cell Signaling Technology, Danvers, MA; 1:1000 dilution), α-mCherry (cat no. 43590S, Cell Signaling Technology, 1:1000 dilution), α-DYKDDDDK tag (cat no. 14793S, Cell Signaling Technology, 1:1000 dilution) for EYA1, AKAP11, and TSHR, α-RET (cat no. 14556, Cell Signaling Technology, 1:1000), α-GAPDH (cat no. 97166S, Cell Signaling Technology, 1:5,000 dilution), α-beta-actin (cat no. 3700S, Cell Signaling Technology, 1:5000 dilution) and α-alpha-tubulin (cat no. MS581P1, Millipore Sigma, 1:5000 dilution). After primary antibody incubation, the membrane was washed three times in 1x TBS-T (cat no. 97062-370, VWR, Radnor, PA) containing 0.1% Tween 20 (P7949, Millipore Sigma) (60 rpm, 5 min, room temperature) and incubated in secondary antibody mixture (60 rpm, 1 h, room temperature). All secondary antibodies were diluted in 1:1 Intercept (LI-COR):1x TBS-T. Secondary antibodies (LI-COR; 1:15,000 dilution) used were: 680LT donkey α-mouse (cat no. 926-68022), 680LT donkey α-rabbit (cat no. 926-68023), 800CW donkey-anti-mouse (cat no. 926-32212), and 800CW donkey α-rabbit (cat no. 925-32213). After secondary antibody incubation, the membrane was washed three times in 1x TBS-T (60 rpm, 5 min, room temperature) and then scanned on an Odyssey CLx Imaging System (LI-COR). At least two independent experiments were performed on different days for each condition.

### Western blot quantification

Protein band intensities were quantified using FIJI (ImageJ)^92^. For each blot, brightness and contrast were adjusted uniformly across the entire image to improve visualization and facilitate consistent band selection; no local or lane-specific adjustments were applied, and care was taken to avoid signal saturation or clipping. Identical horizontally oriented rectangular regions of interest were then placed across all lanes to measure the integrated intensity of bands corresponding to the overexpressed protein of interest (POI), the loading control protein, and mCherry. POI and mCherry intensities were first normalized to the corresponding loading control to account for differences in sample loading. POIs were subsequently normalized to mCherry signal to control for variation in transfection efficiency. We used a paired t-test to test for differences in protein expression between the human and LCA alleles of each protein.

### Investigation of relationship between conservation and effect on chromatin accessibility

For this analysis, when we had sufficient statistical power we generally focused on substitutions in sites with PhyloP > 6 (termed highly conserved sites). If we lacked sufficient power, we instead focused on sites with PhyloP > 3 (termed conserved sites). To test whether substitutions in highly conserved sites were more likely to decrease CA, we used the binomial test comparing the number of human-derived substitutions in highly conserved sites with predicted log_2_ fold-change in CA < -0.5 to the number with predicted log_2_ fold-change in CA > 0.5 where the log_2_ fold-change was always computed with the derived allele in the numerator and the ancestral allele in the denominator. We repeated this analysis for substitutions with absolute predicted log_2_ fold-change in CA in the range 0.25-0.5.

To experimentally validate our findings, we intersected the substitutions in conserved sites with the ATAC-seq peaks with significant differences in allele-specific chromatin accessibility in glutamatergic neurons^48^. We then restricted to only human-derived substitutions with predicted log_2_ fold-change in CA < -0.5 and used Fisher’s exact test to test whether there was an enrichment of peaks with a bias toward lower CA from the human allele in the neuron data. To investigate potential effects on expression, we restricted to only ATAC-seq peaks near genes with ASE (FDR < 0.05) and then repeated this analysis using allele-specific expression instead of allele-specific CA. There was no filtering on whether substitutions were adjacent to HAGs or CA-HAGs.

Next, we investigated the relationship between conservation and total accessibility. For each cell type, we sorted the filtered human-derived substitutions by the maximum of their predicted CA between the human and chimpanzee alleles and took the top 10% most accessible as candidate CREs. We then tested whether highly conserved (PhyloP > 6) sites were enriched in that top 10% using Fisher’s exact test.

### Testing for reductions in evolutionary constraint

To test for differences in the number and conservation of common single nucleotide polymorphisms (SNPs) near fixed substitutions in conserved sites with predicted effects on CA, we used our previously published set of human SNPs with minor allele frequency in the range 0.25-0.75^45^. We only used SNPs within 100 bases of a qualifying fixed substitution. We used independent t-tests to test for differences in mean PhyloP and Fisher’s exact tests for differences in the number of SNPs relative to that expected based on the number of fixed substitutions. To test for differences in ASE variance, we used our previously published measurements of ASE variance derived from data produced by the GTEx Consortium^93,94^, filtered to remove the very small number of outlier genes with ASE variance > 0.5 and any genes with fewer than 100 individuals with computable ASE. For human-chimpanzee ASE, we used previously published data from nine cellular contexts: cardiomyocytes, skeletal muscle, hepatic progenitors, pancreatic progenitors, retinal pigmented epithelial cells, cortical organoids after 50 days of differentiation, cortical organoids after 100 days of differentiation, and cranial neural crest cells^26,27,95^.

To estimate within-human ASE variance in relevant tissues, for cardiomyocytes we used heart data from GTEx, for cortical organoids and motor neurons we used brain data, for hepatic progenitors we used liver, and for all other cell types we pooled data across all GTEx tissues. As genes with lower expression from the human allele generally had fewer individuals with sufficient reads to estimate allele-specific expression in the GTEx data, we down-sampled the genes with higher expression from the human allele to match the distribution of the number of individuals with measurable ASE variance for the genes with lower expression from the human allele. We repeated this procedure 100 times, using the Mann-Whitney U test to test for a difference in the median ASE variance. We report the median p-value across the 100 down-samplings for each cell type.

### Analysis of recent selection on conserved sites

We downloaded the publicly available selection summary statistics from Akbari et al.^50^, lifted over to Hg38, and intersected them with our PhyloP and nearest gene annotations^96^. We flipped the sign of the selection coefficient (S) when the ancestral allele matched the alternate allele so that it always represented the strength of selection on the derived allele. For the relaxed cutoff, we considered anything with nominal p-value < 10^-5^ to be selected and for the strict cutoff we required FDR < 0.05 (equivalent to p-value < 5.09x10^-7^). We binned sites by PhyloP score and tested for enrichment of selected sites (defined as above) against sites with no evidence for selection (p > 0.05 and |S| < 0.001). For nonsynonymous substitutions, we intersected with our previously generated list of polymorphic nonsynonymous substitutions^45^, split sites into strongly selected (|S| > 0.015) and weakly selected (p-value or FDR as described above and |S| < 0.015), and used the Mann-Whitney U test to test for differences in the medians of the PhyloP distributions.

### GWAS Catalog Annotation

To annotate these sites with known phenotypic associations, each candidate position was first mapped to a dbSNP rsID using the Ensembl REST API overlap endpoint^97^. Where multiple variants were reported at the same position, allele-matching against the known REF and ALT alleles was preferred; otherwise the first returned rsID was retained. The GWAS Catalog^51^ was filtered to genome-wide significant associations (p ≤ 5×10⁻) prior to annotation. For each candidate variant, the rsID of the input site and all linkage disequilibrium (LD) proxy rsIDs (see below) were matched against the “SNPS” field of the filtered GWAS Catalog. This procedure produced a table of input-position × GWAS-trait pairs that formed the basis of downstream analyses. Each unique GWAS trait string was classified into two levels of ontological granularity (broad and fine) using a keyword-based classifier informed by the Experimental Factor Ontology (EFO)^52^.

### Linkage Disequilibrium Proxy Identification

To capture GWAS associations at variants in LD with each candidate site, we used per-chromosome high-coverage phased VCF files from the 1000 Genomes Project GRCh38 release^58^, obtained from the International Genome Sample Resource. For each autosome, a BED file of candidate positions was constructed and plink2^98^ was used to compute pairwise LD with the flags --r2-unphased --ld-window-kb 500 --ld-window-r2 0.8, restricting to EUR samples. The resulting LD pairs were assembled into a proxy lookup table. To support signed-correlation calculations required for derived-allele direction assignment (see below), plink2 was additionally run with --r-unphased to obtain the signed Pearson correlation coefficient (r) for each proxy pair. The query variant itself was always included as a direct hit. In total, each candidate site was annotated with all proxy rsIDs satisfying R² ≥ 0.8 in the EUR population within a ±500 kb window.

### Derived Allele Effect Direction

For each input-position × GWAS-trait pair, the direction of the derived allele’s effect on the trait was inferred in a multi-step procedure. First, the risk allele was parsed from the “STRONGEST SNP-RISK ALLELE” field of the GWAS Catalog, which encodes alleles in the format rsXXXXXX-A (rsID followed by a dash and the risk allele base). Second, the direction of the risk allele’s effect was determined from the reported effect estimate: associations with OR or Beta > 1, or whose 95% confidence interval text contained the word “increase,” were classified as alternative-allele-increasing; those with OR or Beta < 1, or CI text containing “decrease,” were classified as alternative-allele-decreasing. All other categories were removed. We then restricted only to “direct hits” and sign of the effect was flipped if the derived allele from the Akbari et al. table did not match the alternative allele from the GWAS catalog and labeled appropriately.

### Integrative analysis of recent selection, conservation, GWAS, and effects on CA

Analyses were carried out for selection for and against the derived allele separately. To test which broad GWAS categories were most enriched for substitutions in highly conserved (PhyloP > 6) sites, we used a one-sided Fisher’s exact test to test whether substitutions in each category (with duplicate GWAS tag SNPs and substitutions removed) were enriched for substitutions in highly conserved sites relative to all substitutions associated with at least one GWAS association (with duplicate GWAS tag SNPs and substitutions removed separately in this background set) for sites with. We repeated this approach for the finer GWAS categories within broad categories that had FDR < 0.05. For each trait in at least one FDR significant fine category, we tested whether there was a bias toward the derived allele increasing or decreasing the trait using Fisher’s exact test.

Next, we intersected the list of sites with our previously published set of predicted effects of human polymorphisms on CA in fetal cortical excitatory neurons^45^. We restricted to conserved sites (PhyloP > 3) and then used a one-sided Fisher’s exact test to test whether substitutions associated with cognitive or psychiatric traits were enriched for decreases (predicted log_2_ fold-change in CA < -0.25) or increases (predicted log_2_ fold-change in CA > 0.25) in CA relative to substitutions not associated with a cognitive or psychiatric trait. We also used a one-sided Fisher’s exact test to determine whether positively selected substitutions associated with cognitive or psychiatric traits were enriched for predicted decreases in CA relative to negatively selected substitutions. The plot of rs1334297 derived allele frequency over time was created at https://reich-ages.rc.hms.harvard.edu/.

## Statistics

We note the statistical tests used throughout the methods section and results. All statistical tests, with the exception of mashr, were implemented in scipy v1.15.3^99^. All tests are two-sided unless stated otherwise.

## Acknowledgements

Biorender was used in the creation of some figure panels.

## Author Contributions

A.L.S. conceived the project. A.L.S. performed all analysis, visualization, validation, and writing of software with guidance from H.B.F. A.L.S. and G.M.C. did the subcloning. L.M. and G.M.C. did all cell culture, Western blots, and Western blot quantification. M.E.P. did the ChromBPNet model training and inference. A.L.S., G.M.C., L.M., and H.B.F. wrote the article and A.L.S. created the figures with input from H.B.F. H.B.F. provided funding for the study. All authors approved the final version of the manuscript.

## Data and code availability

The data from the study of human-chimpanzee hybrid cortical organoids are available from the Gene Expression Omnibus (GEO) accession GSE144825. The hybrid expression data from six cell types are available from GEO with accession GSE232949. Hybrid expression data from CNCCs are available from GEO with accession GSE146481. ChromBPNet model weights are deposited at: http://mepalmer.net/34_models.tgz. Ancient DNA-based quantification of natural selection is available at: https://dataverse.harvard.edu/dataset.xhtml?persistentId=doi:10.7910/DVN/7RVV9N. Thousand genomes VCFs are available from: https://ftp.1000genomes.ebi.ac.uk/vol1/ftp/data_collections/1000G_2504_high_coverage/. Developing human neocortex single cell gene expression profiles are available from: https://datasets.cellxgene.cziscience.com/7b582c37-37ad-49d2-a369-06f38289031e.h5ad. Code is available at: https://github.com/astarr97/AcceleratedEvolution.

## Description of supplemental files

**Supplemental Figures:** Contains supplemental figures 1-9.

**Supplemental Table 1:** Information for all proteins tested for accelerated evolution.

**Supplemental Table 2:** Information for all gene ontology biological process (GOBP) and human phenotype (HPO) terms tested for accelerated protein evolution.

**Supplemental Table 3:** Information for all genes tested for 3’ UTR and 5’ UTR accelerated evolution.

**Supplemental Table 4:** Information for all gene ontology biological process (GOBP) tested for accelerated 5’ UTR evolution.

**Supplemental Table 5:** Information for all genes tested for accelerated evolution of their *cis*-regulatory neighborhoods.

**Supplemental Table 6:** Information for all gene ontology biological process (GOBP) tested for accelerated *cis*-regulatory neighborhood evolution.

**Supplemental Table 7:** Per 500 base pair window difference in PhyloP test statistics for human-accelerated genes to prioritize individual regions for follow-up.

**Supplemental Table 8:** Information for all genes tested for accelerated evolution of chromatin accessibility across 34 cell types.

**Supplemental Table 9:** Predicted effects on CA and total accessibility across 34 cell types for all human-derived substitutions in conserved sites (PhyloP > 3).

**Supplemental Zip File 1:** Annotated plasmid sequences used in this study.

**Supplemental Text 1:** Discussion of alternative explanations for our observations of accelerated evolution.

**Supplemental Text 2:** Discussion comparing our results to those previously obtained for protein-coding regions using dN/dS-like methods.

**Supplemental Text 3:** Discussion of global shift toward substitutions in conserved sites in the human lineage and the reasoning behind controlling for this global shift.

**Supplemental Text 4:** Discussion of the relationship between conservation and recent positive selection on nonsynonymous variants in human evolution.

**Supplemental Text 5:** Discussion of limitations of cognitive and behavioral GWAS and how it relates to our results.

