## Supplemental Figures for "Adaptive loss of function accelerated the evolution of ancient and modern human cognition"

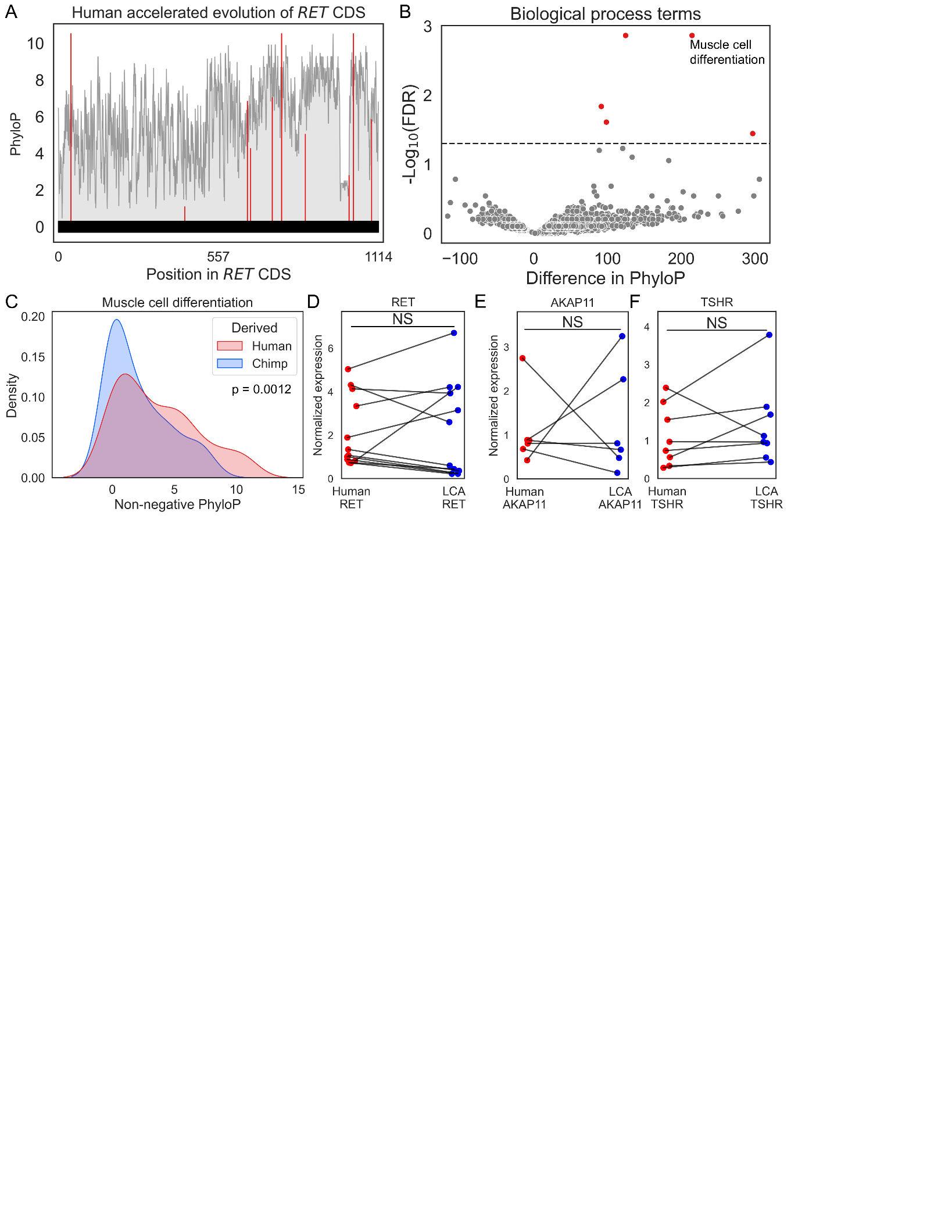

**Supplemental Figure 1: A)** Plot showing PhyloP scores in the *RET* CDS. Red indicates sites with human-derived substitutions. The grey trace is the moving average of PhyloP scores across all sites. **B)** Protein acceleration for different GO biological process terms. **C)** PhyloP distributions for human-derived (red) and chimpanzee-derived (blue) nonsynonymous substitutions for proteins associated with muscle cell differentiation. **D-F)** Normalized protein expression for human and human-chimpanzee LCA RET **(D)**, AKAP11-FLAG **(E)**, and TSHR-FLAG (**F)**.

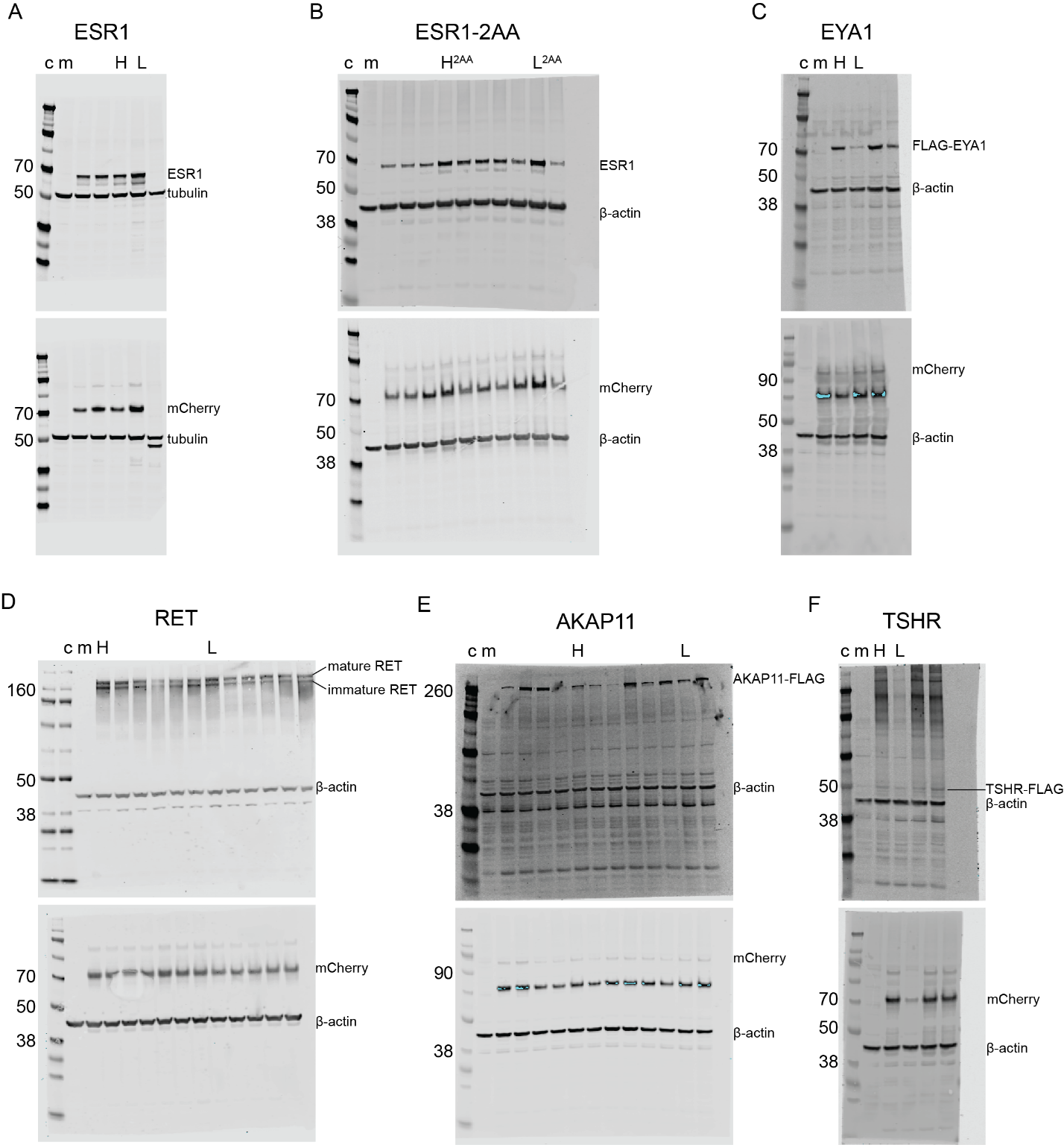

**Supplemental Figure 2:** Western blots showing protein expression of ESR1 **(A)**, ESR1 with residues 300 and 306 swapped between human and LCA (2AA) **(B)**, FLAG-EYA1 **(C)**, RET **(D)**, AKAP-FLAG **(E),** and TSHR-FLAG **(F)**. c: Chameleon Duo Pre-Stained Marker, m: mock transfected, H: human, L: last common ancestor. Only representative lanes are annotated; uncropped immunoblots are shown for completeness. The same lysates were run in duplicate (top and bottom panels) and then immunoblotted for the indicated proteins. At least two independent biological replicates were performed for each condition.

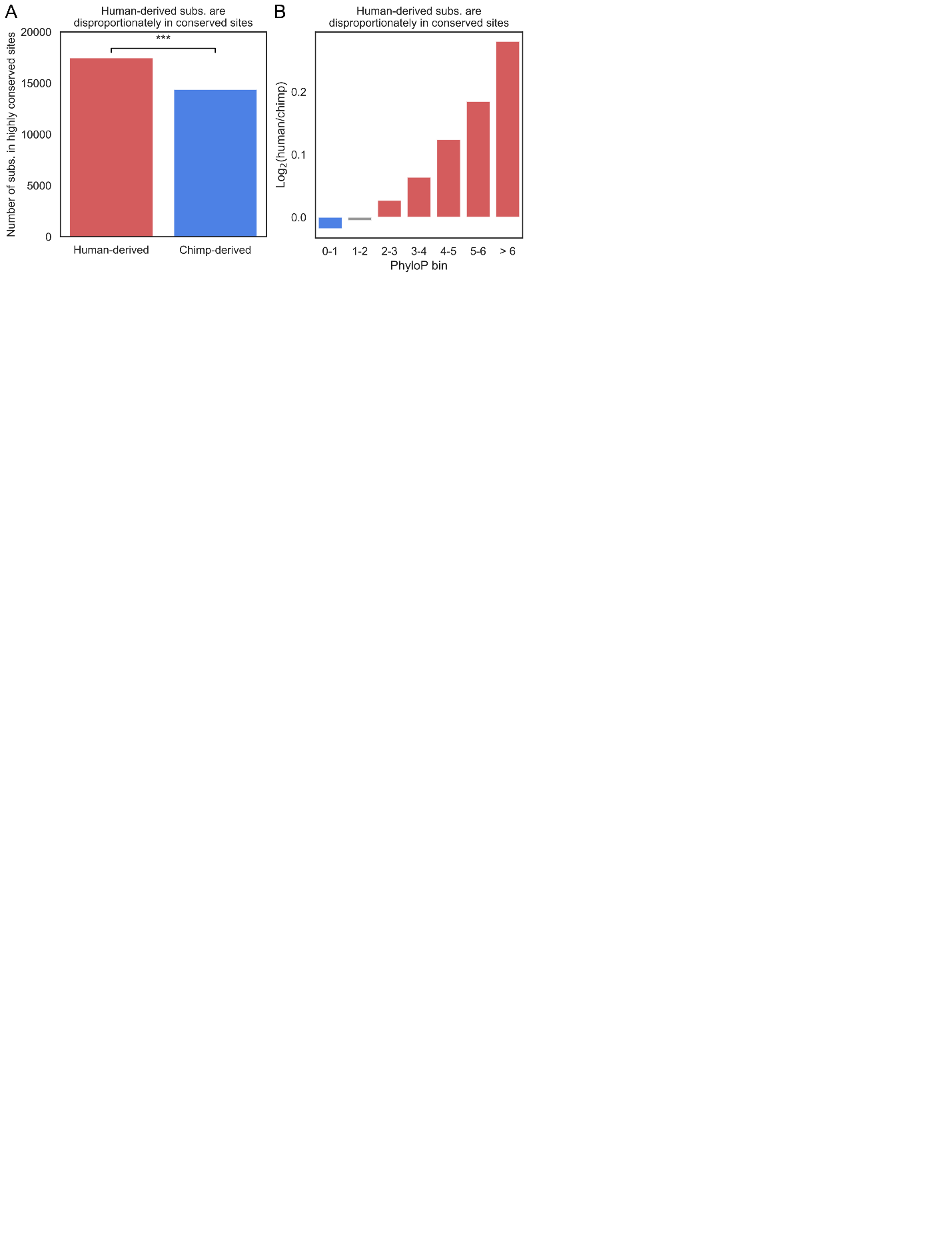

**Supplemental Figure 3: A)** Number of human-derived and chimpanzee-derived substitutions in highly conserved sites (PhyloP > 6). *** p < 0.0005. **B)** Enrichment of human-derived substitutions relative to chimpanzee-derived (y-axis) in increasing PhyloP bins (x-axis).

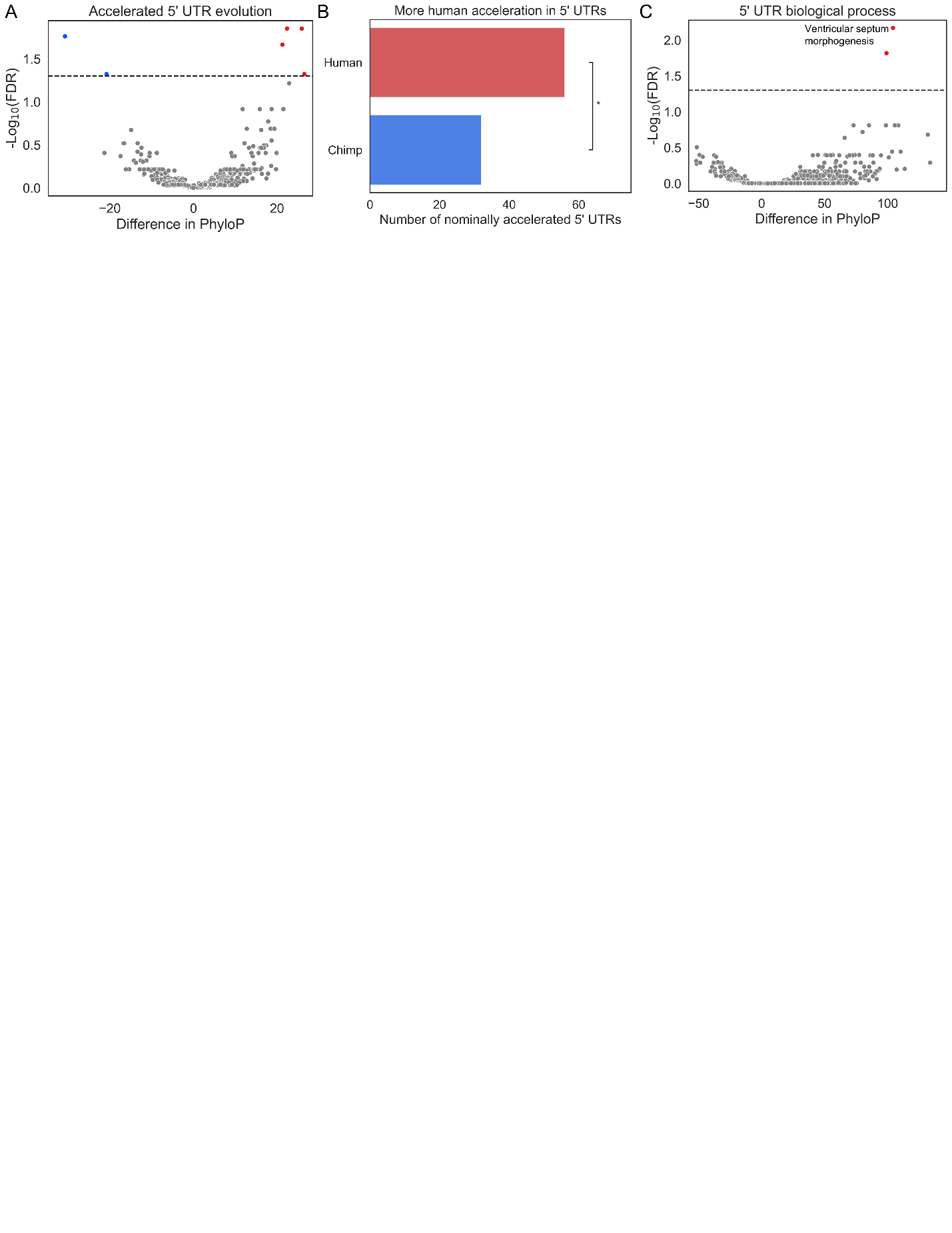

**Supplemental Figure 4: A)** 5’ UTR acceleration. Human-accelerated 5’ UTRs are shown in red and chimpanzee-accelerated in blue. **B)** Number of nominally human-accelerated and chimpanzee-accelerated 3’ UTRs. * p < 0.05. **C)** Same as in (A) but for GO biological process terms.

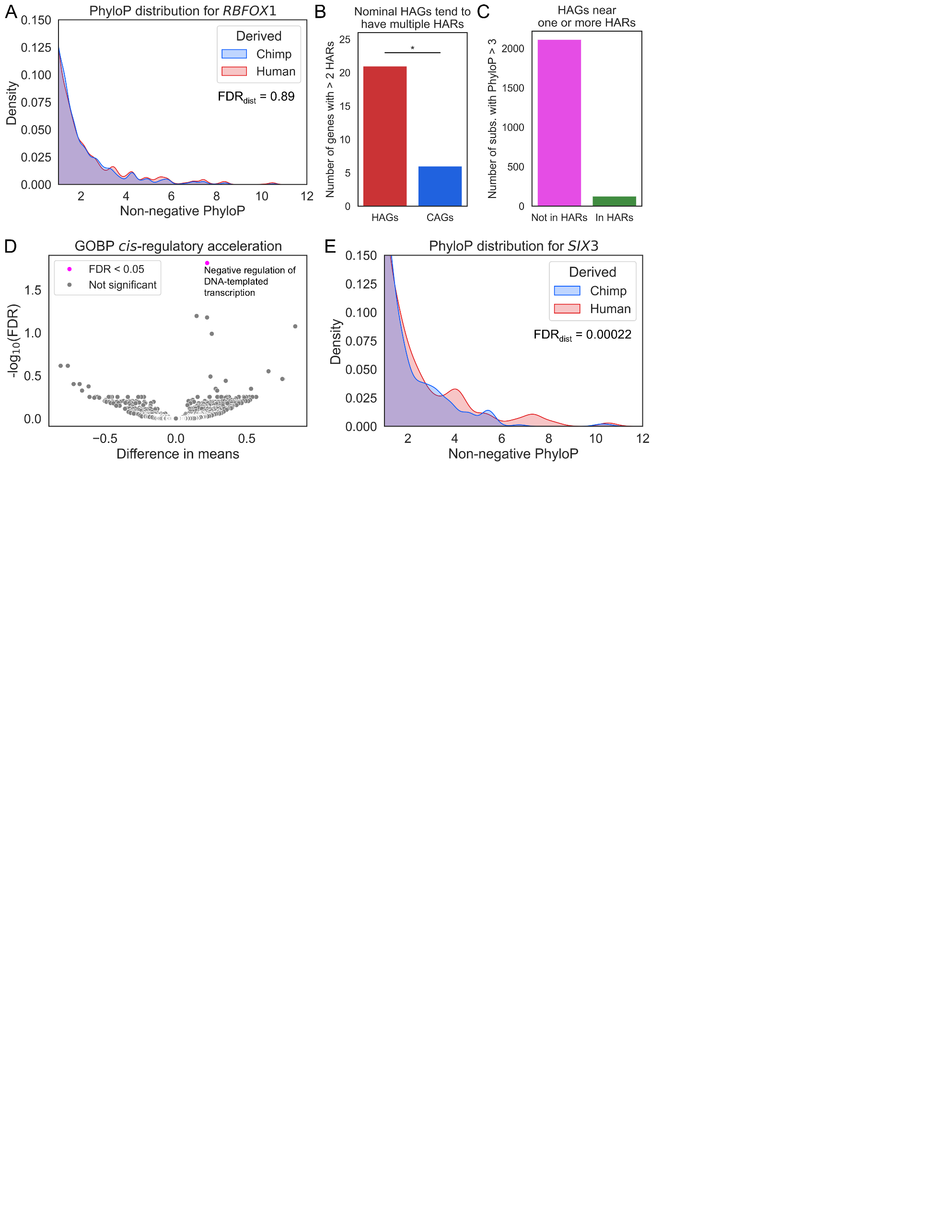

**Supplemental Figure 5: A)** Distribution of PhyloP scores for substitutions in the *cis*-regulatory neighborhood of *RBFOX1*. **B)** Number of human-accelerated genes (HAGs) with at least three nearby HARs compared to the number of chimpanzee-accelerated genes (CAGs). * p < 0.05. **C)** Number of conserved sites near HAGs with at least one nearby HAR that are (green) or are not (magenta) in HARs. **D)** Volcano plot from comparing acceleration z-scores for a GO BP category against all genes not in that category. P-values are from independent t-test. **E)** Same as in (A) except for *SIX3*.

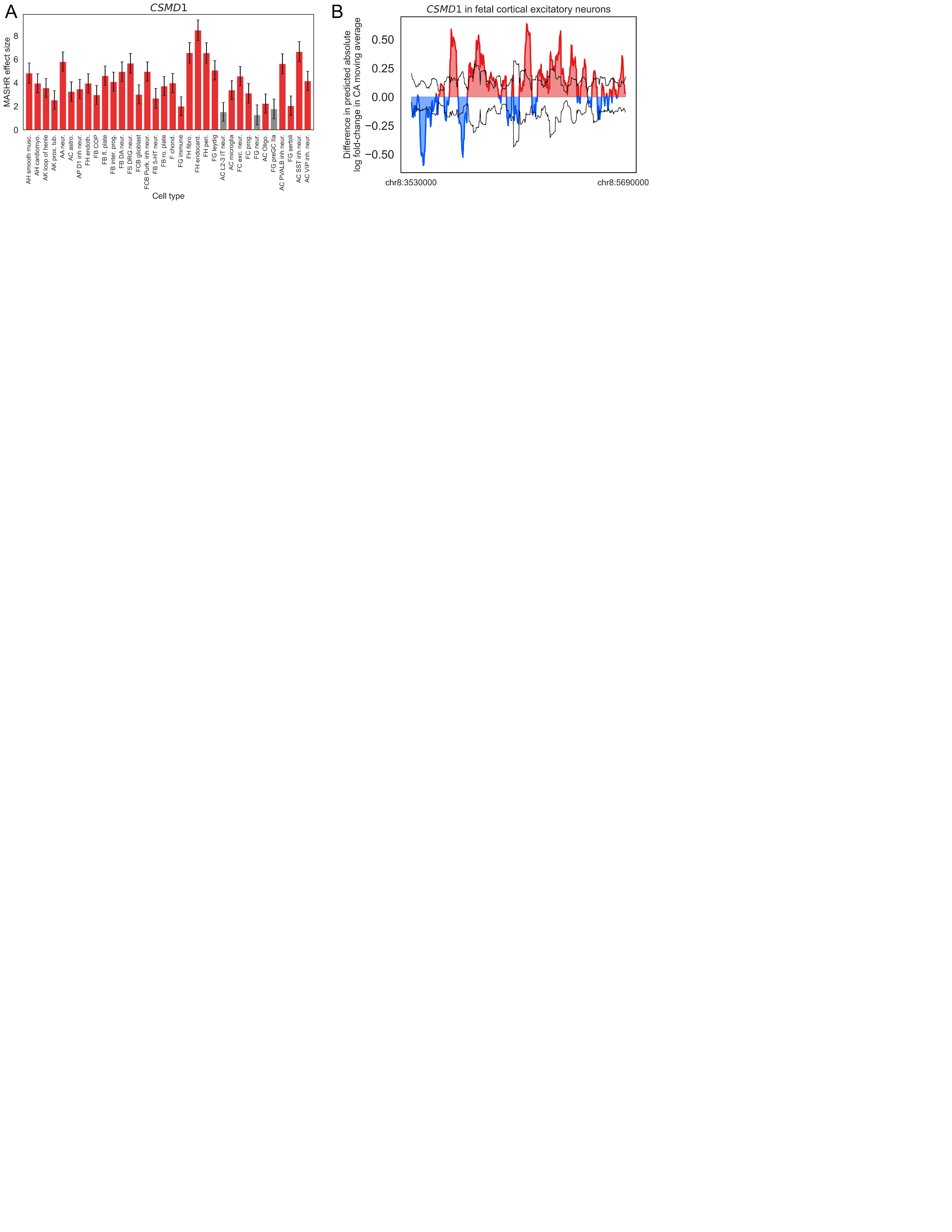

**Supplemental Figure 6: A)** Estimated effect sizes from mashr for accelerated evolution of CA in each cell type for *CSMD1*. Error bars are the posterior standard deviations. Cell types with significantly human-accelerated CA evolution are shown in red. **B)** Accelerated evolution of CA for *CSMD1* in humans. Colored, filled trace shows the moving average of the difference in the sum of human-derived and chimpanzee-derived predicted absolute log_2_ fold-changes. The black line indicates one standard deviation above and below the mean absolute log_2_ fold-change difference of 100 iterations permuting which lineage each site was assigned to.

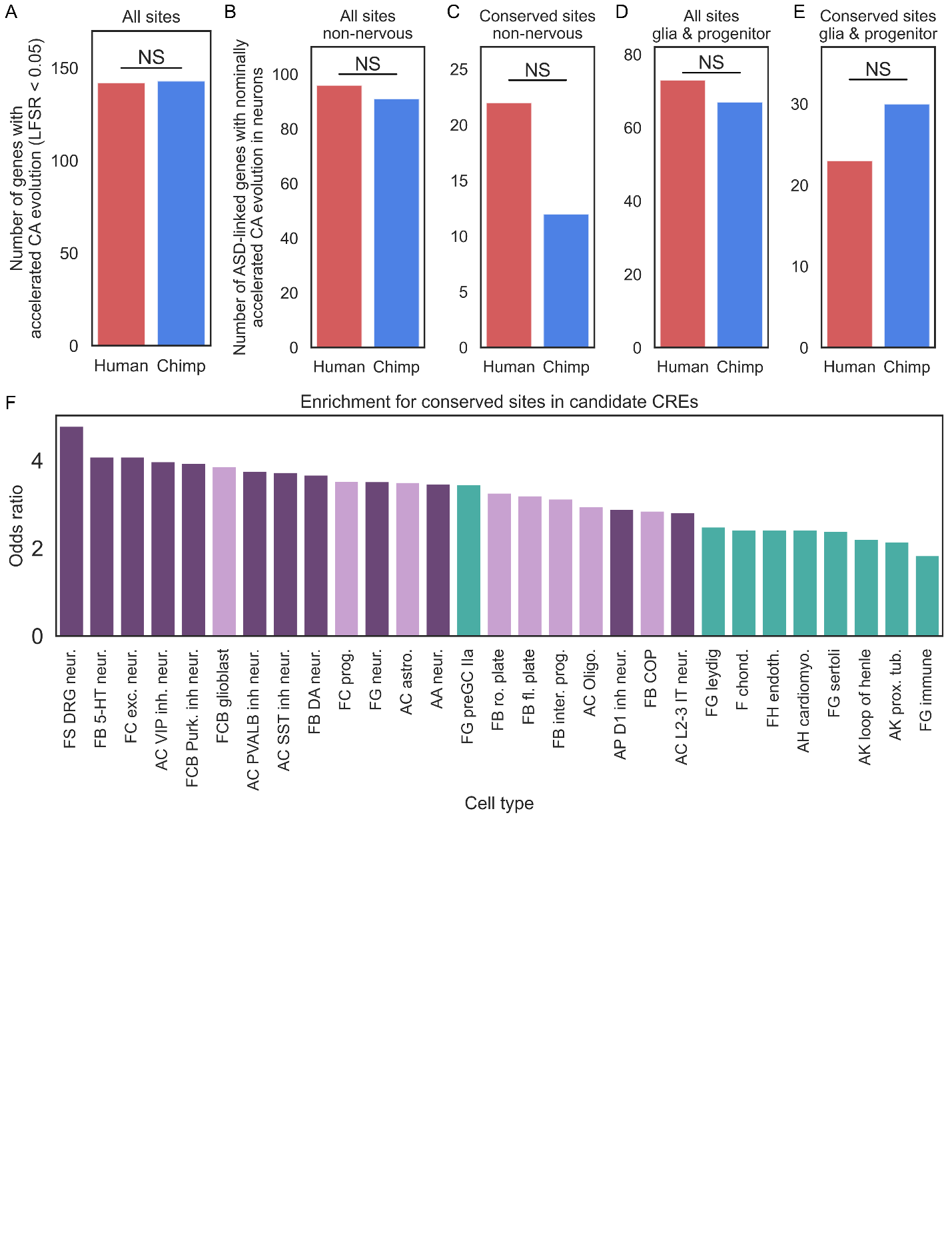

**Supplemental Figure 7: A)** Number of genes with significantly accelerated evolution of CA (mashr local false sign rate < 0.05 in at least one cell type) using all sites in the top decile of predicted CA as input. **B)** Number of ASD-linked genes with nominally accelerated evolution of CA in at least one non-neuronal cell type. **C)** Same as in (B) but restricting to sites with PhyloP > 1. **D)** Same as in (A) but for glia and progenitor cells. **E)** Same as in (D) but restricting to sites with PhyloP > 1. **F)** Enrichments for human-derived substitutions in conserved sites in the top decile of accessibility (same as in Fig. 4H but with cell type labels added).

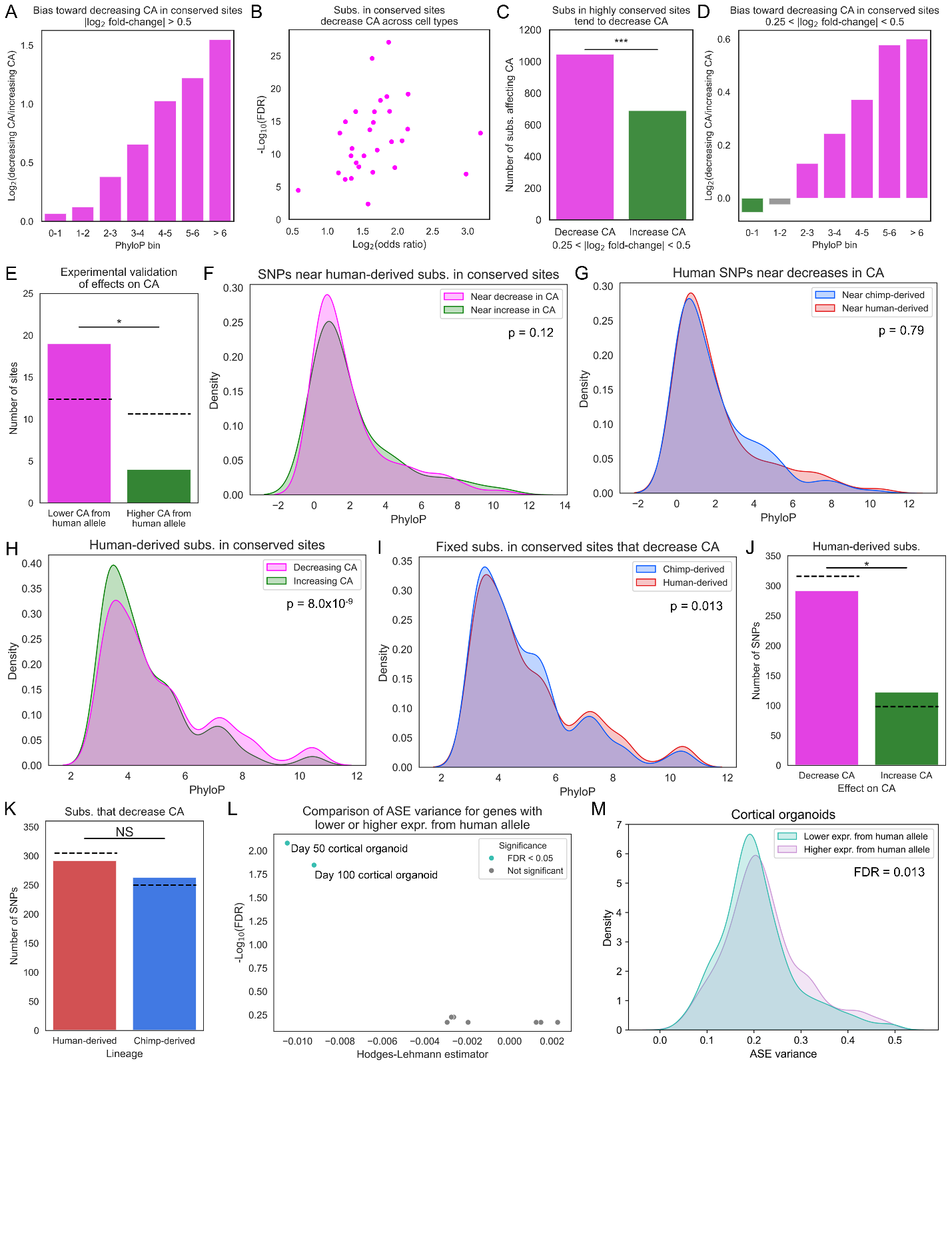

**Supplemental Figure 8: A)** Enrichment of substitutions that decrease CA over sites that increase CA in increasingly conserved sites. **B)** Enrichment for substitutions in highly conserved sites being predicted to decrease CA (predicted log_2_ fold-change in CA < -0.5) relative to increase CA across cell types. All FDR < 0.05. Odds ratio reflects the enrichment for highly conserved sites relative to those with PhyloP in the range 0-1. **C)** Bar height shows the number of substitutions in highly conserved sites (PhyloP > 6) that are predicted to decrease CA (-0.5 < predicted log_2_ fold-change < -0.25) or increase CA (0.25 < predicted log_2_ fold-change < 0.50). *** p < 0.0005. **D)** Same as in (A) but for 0.25 < |predicted log_2_ fold-change| < 0.5. **E)** Number of substitutions in highly conserved sites (PhyloP > 6) with predicted log fold-change < -0.5 that are in ATAC peaks with lower accessibility from the human allele (magenta) and higher accessibility from the human allele (green). Dashed lines indicate the number expected by chance. * p < 0.05. **F)** PhyloP distribution for polymorphisms near fixed, conserved (PhyloP > 3), human-derived substitutions that decrease CA (magenta) or increase CA (green). Only substitutions predicted to have an absolute log_2_ fold-change in CA > 0.5 were included. P-value is from t-test. **G)** Same as in (D) but for polymorphisms near chimpanzee-derived (blue) and human-derived (red) substitutions predicted to decrease CA. **H)** PhyloP distributions for fixed, human-derived substitutions in conserved sites (PhyloP > 3) for those predicted to decrease CA (magenta) and increase CA (green). **I)** Same as in (F) but for human-derived (red) and chimpanzee-derived (blue) substitutions that are predicted to decrease CA. **J)** Bar height shows the number of common human SNPs near fixed human substitutions in conserved (PhyloP > 3) sites that are predicted to decrease CA (magenta) or increase CA (green). Dashed line shows the number expected by chance. * p = 0.05. **K)** Same as in (H) but for SNPs near human-derived (red) or chimpanzee-derived (blue) substitutions that are predicted to decrease CA. **L)** Volcano plot from using the Mann-Whitney U test to determine whether genes with lower expression from the human allele have significantly higher or lower ASE variance than genes with higher expression from the human allele across nine cell types. **M)** Distribution of ASE variance for genes with lower expression from the human allele in day 100 cortical organoids (teal) and higher expression from the human allele (purple). FDR is from Benjamini-Hochberg corrected Mann-Whitney U test p-value.

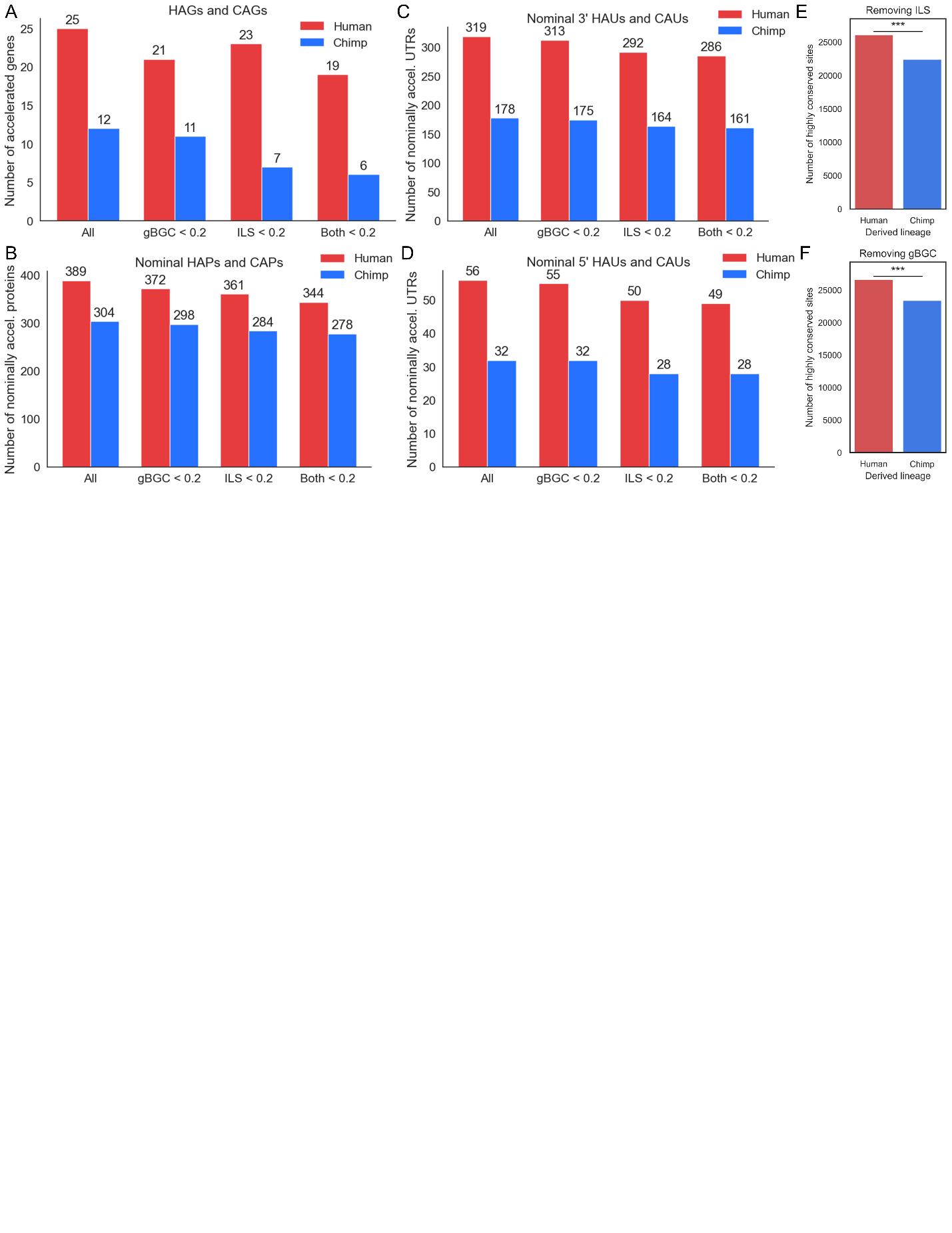

**Supplemental Figure 9**: **A)** Number of HAGs (red) and CAGs (blue) restricting to, from left to right: all accelerated genes (AGs), AGs with fraction of signal explained by GC-biased gene conversion (gBGC) < 0.2, AGs with fraction of signal explained by incomplete lineage sorting (ILS) < 0.2, both < 0.2. **B)** Same as in (A) but for nominal protein acceleration. **C)** Same as in (A) but for nominal 3’ UTR acceleration. **D)** Same as in (A) but for nominal 5’ UTR acceleration. **E)** Comparison of number of human- and chimp-derived substitutions in highly conserved sites removing ILS sites. **F)** Comparison of number of human- and chimp-derived substitutions in highly conserved sites removing gBGC sites. *** indicates p < 0.0001.

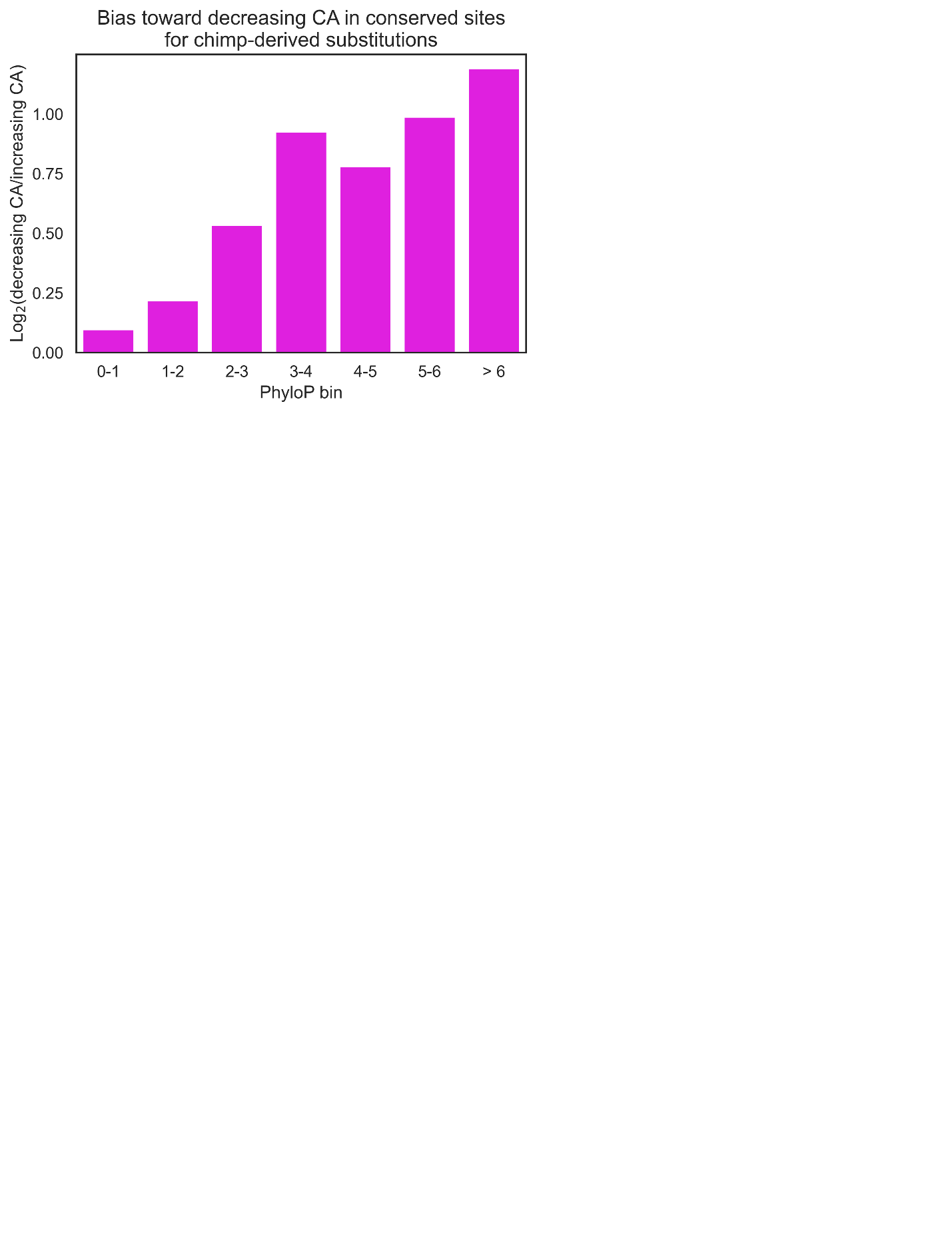

**Supplemental Figure 10**: Enrichment of chimp-derived substitutions that decrease CA over sites that increase CA in increasingly conserved sites.

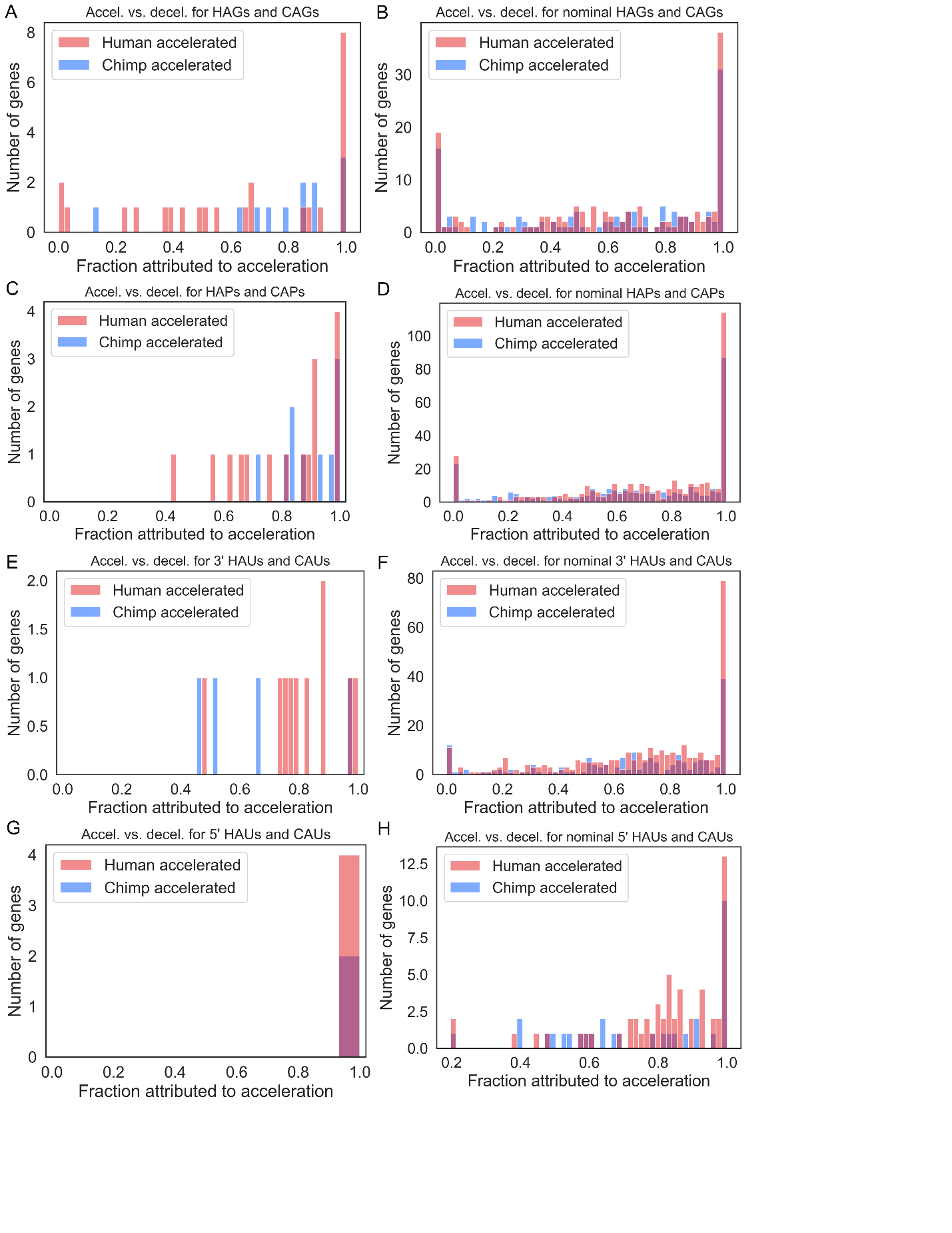

**Supplemental Figure 11: A)** Histogram of fraction of acceleration explainable by acceleration relative to the gorilla lineage across FDR-significant HAGs and CAGs. **B)** Same as in (A) but for nominally including nominally significant HAGs and CAGs. **C-D)** Same as in (A-B) but for proteins. **E-F)** Same as in (A-B) but for 3’ UTRs. **G-H)** Same as in (A-B) but for 5’ UTRs.

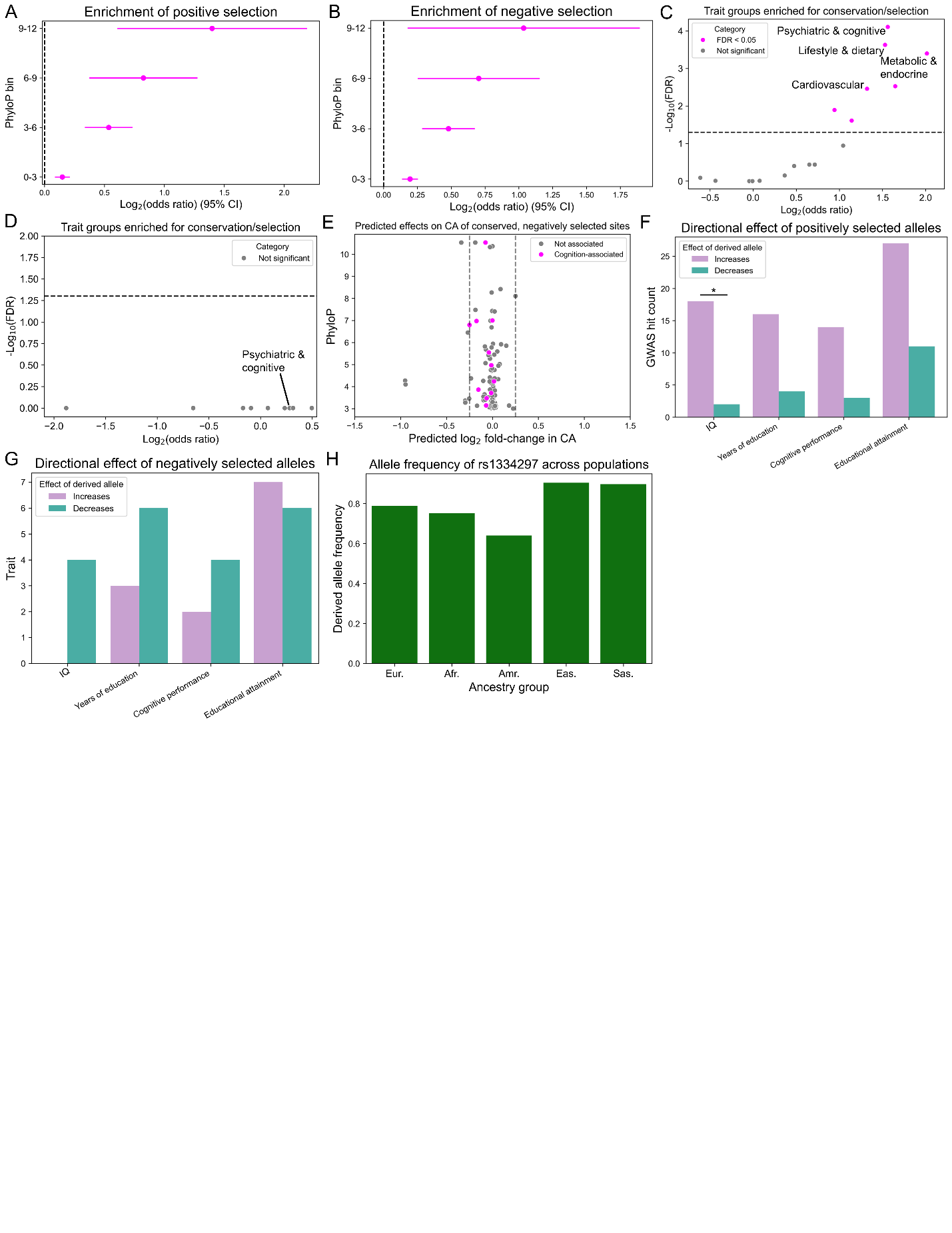

**Supplemental Figure 12: A)** Enrichment of recently positively selected substitutions in conserved sites with strong evidence for positive selection (FDR < 0.05). **B)** Same as in (A) but for negative selection and at relaxed p < 10^-5^ cutoff (to match Fig. 6A). **C)** Positively selected GWAS hit enrichments for highly conserved (PhyloP > 6) sites. Different GWAS were grouped by broad MeSH similarity. **D)** Same as in (C) but for negatively selected substitutions. **E)** PhyloP scores (y-axis) and predicted effects on CA of the derived allele in fetal cortical neurons (x-axis) for sites with negative selection. Sites are magenta if they are associated with cognitive or psychiatric phenotypes. **F)** Directional effects of positively selected alleles that are direct GWAS hits. Only GWAS terms with nominal p < 0.05 are shown. * FDR < 0.05. **G)** Directional effects of negatively selected alleles that are direct GWAS hits. Only GWAS terms from (F) are shown. **H)** Derived allele frequency of chr13:57761241-A across modern human populations.

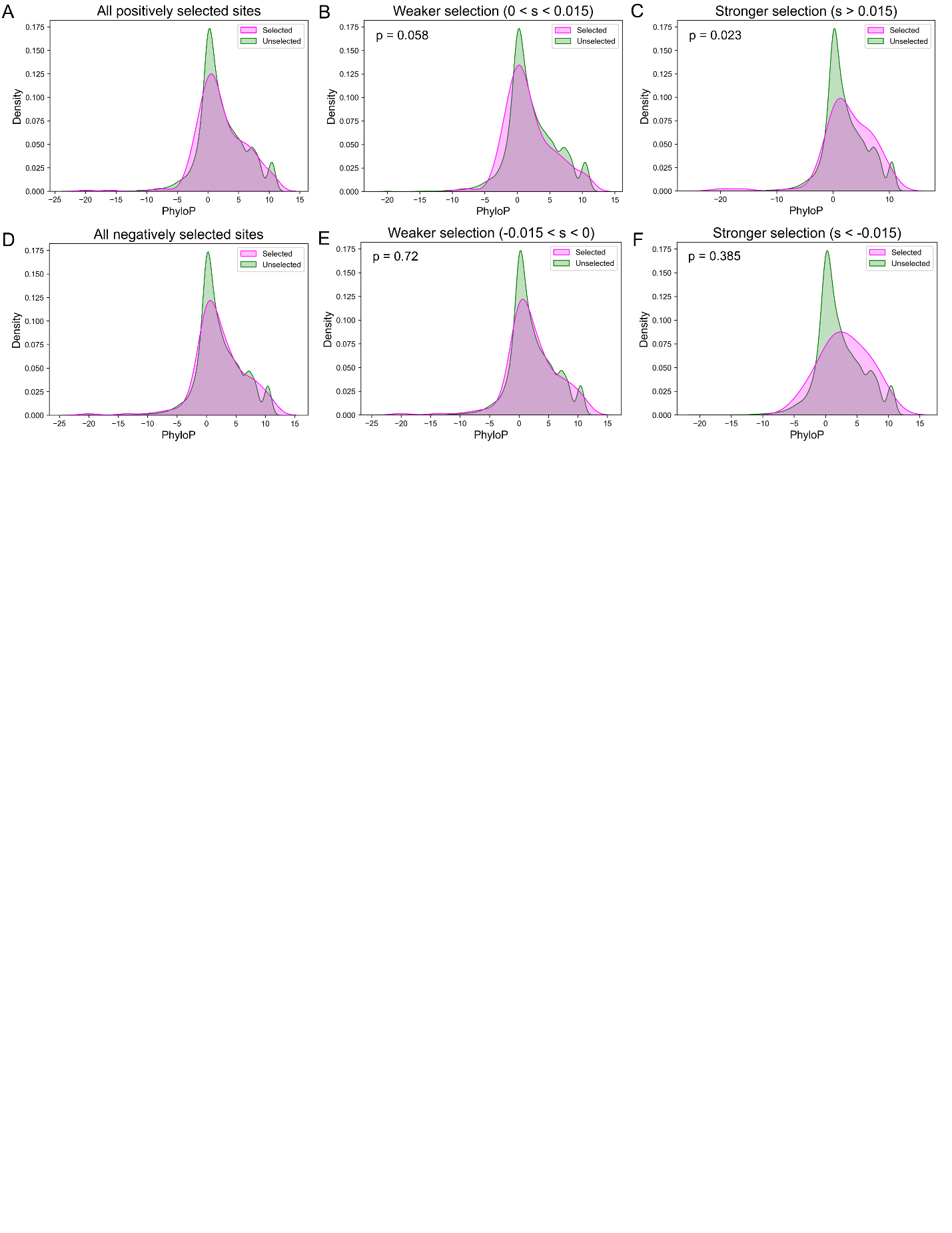

**Supplemental Figure 13: A)** PhyloP distributions for sites with substitutions under positive selection. **B)** Same as in (A) but for weakly positively selected substitutions only. P-value is from Mann-Whitney U test. **C)** Same as in (A) but for strongly positively selected substitutions only. **D)** Same as in (A) but for negatively selected substitutions. **E)** Same as in (A) but for weakly negatively selected substitutions. **F)** Same as in (A) but for strongly negatively selected substitutions.
