## Supplemental Text 1 for "Adaptive loss of function accelerated the evolution of ancient and modern human cognition"

Here, we detail the potential roles of various causes of accelerated evolution beyond positive selection. These include other modes of selection (reduced constraint), mutational processes, and technical factors. Finally, we discuss the potential role of deceleration in one lineage relative to another in driving our results.

**Reductions in evolutionary constraint**

A human-derived substitution causing a large decrease in CA may result from three possible evolutionary scenarios (summarized in Table 1). First, if a range of CA at a particular CRE is selectively neutral in humans and chimpanzees, then the substitution could have drifted to fixation without any role of natural selection. Second, the CRE in question could have lost evolutionary constraint only in the human lineage, allowing the substitution that decreased CA to reach fixation. Third, there may have been positive selection for decreased CA at that CRE, in which case the substitution provided a fitness advantage to ancient humans. Although it is difficult to fully distinguish these three scenarios, our observation of greater *cis*-regulatory acceleration in the human lineage argues against the first scenario.

To test the second hypothesis of broad, human-specific reduced constraint, we analyzed within-human variation^1^. A decrease in constraint is expected to lead to increased within-species polymorphism, as compared to either 1) the same regions in a sister species where constraint has not been lost, or 2) different regions in the same species where constraint has not been lost. Therefore, a major prediction of the loss-of-constraint model is that regions with human-derived substitutions that decreased CA should show greater within-human polymorphism as compared to 1) the same regions in chimp, and 2) regions with human-derived substitutions that increased CA (which are inherently matched for a multitude of potential confounding factors with substitutions that decreased CA). Performing both of these comparisons using the CA predictions from Purkinje cells, we observed no significant difference in mean PhyloP between polymorphisms near human-derived substitutions that decreased CA and those that increased CA (Supp. Fig. 8F, p = 0.12, t-test), nor between the former and their chimpanzee-derived counterparts (Supp. Fig. 8G, p = 0.79, t-test). This is despite the human-derived fixed substitutions that decreased CA occurring in more conserved sites than either those that increased CA (Supp. Fig. 8H, p = 8.0x10^-9^, t-test) or chimpanzee-derived substitutions that decreased CA (Supp. Fig. 8I, p = 0.013, t-test). Moreover, when counting the number of variants rather than their conservation, there was a slight depletion of common polymorphisms near the human-derived substitutions that decreased CA relative to those that increased CA (Supp. Fig. 8J, p = 0.05, Fisher’s exact test) and no significant difference when comparing to the chimpanzee-derived substitutions (Supp. Fig. 8K, p = 0.33, Fisher’s exact test). Collectively, this argues against the hypothesis that these substitutions resulted from a decrease in evolutionary constraint, though we cannot entirely rule out this explanation.

Next, we wanted to test whether decreases in gene expression, which are frequently a consequence of decreases in CA (e.g., Fig. 5C-D), might result from decreased constraint. Applying similar logic to that outlined above, we used ASE variance, a previously published measure that quantifies within-human variability in *cis*-regulation (and thus constraint on gene expression during recent human evolution)^2^, with higher ASE variance corresponding to lower constraint. Across nine diverse cell types/organoids differentiated from human-chimpanzee hybrid iPSCs^3–5^, we never observed significantly greater ASE variance for genes with lower expression from the human allele when compared to those with higher expression (while controlling for differences power to estimate ASE variance, Supp. Fig. 8L). In one case, we observed the opposite—lower ASE variance for genes with lower expression from the human allele (cortical organoids FDR = 0.013, Supp. Fig. 8M)—suggesting these genes may be under greater expression constraint in human populations. Although it is difficult to fully rule out all other possible explanations for our observations, these results, combined with disproportionate accelerated evolution in the human lineage and our findings for modern human evolution, suggest that a considerable fraction of decreases in CA caused by human-derived substitutions may have been the result of positive selection (Table 1).

Overall, we find no evidence for loss of constraint near substitutions in conserved sites that are predicted to decrease CA nor for genes with lower expression in humans than chimpanzees. Although it is likely that loss of constraint plays an important role for some fraction of substitutions in conserved sites in humans, combined with evidence from recent human evolution (Fig. 6), this suggests that a considerable portion of the accelerated evolution in conserved sites in the human lineage is explained by positive selection.

**Incomplete lineage sorting**

Here, we use incomplete lineage sorting (ILS) to refer specifically to sites that support tree topologies that are not humans being more closely related to chimpanzees than gorillas (i.e. HCG). For example, we define CGH sites as those where chimpanzee and gorilla have the same base, human has a different base, yet orangutan has the same base as human (and *vice versa* for HGC sites)^6^. Although this could be caused by recurrent substitutions, a more likely explanation is ILS. In principle, a pattern of more CGH sites or more HGC sites could produce signals we defined here as human or chimpanzee acceleration. To understand the role that this might have played in our results, we quantified the sum of the number of CGH and HGC sites, weighted by the non-negative PhyloP scores (the same statistic used to detect accelerated sequence evolution). We did this for protein, UTR, and non-coding region acceleration.

Across all analyses, the proportion of the signal explainable by ILS was less than 0.05 for most genes (Supp. Tables 1, 3, and 5). For example, for HAGs with FDR < 0.05, 17 out of 25 has a proportion less than 0.05. There were a few outlier genes for which ILS explained a large proportion of the signal. For example, for statistically significant HAGs, two genes (*TMPRSS11F* and *FREM3*) had greater than 20% of signal explained by ILS, whereas for CAGs five genes did (*HERC4*, *HOXC4*, *NFYB***,** *POLR3B*, and *GABRR2*) (Supp. Fig. 9A-D).

The other major question is whether ILS alone could be driving the overall bias toward human acceleration that we observed. If CGH sites tend to have much higher PhyloP scores than HGC sites, then this could lead to the genome-wide bias toward higher PhyloP scores for human-derived substitutions that we observed. However, even when removing all CGH and HGC sites, we observed a similar bias toward human-derived substitutions occurring in highly conserved sites more frequently than chimpanzee-derived sits (Supp. Fig. 9E, p = 4.4x10^-82^, binomial test). Overall, this suggests that while ILS may slightly affect the identification of HAGs and CAGs, it cannot explain the genome-wide bias toward human acceleration we observed.

**GC-biased gene conversion**

Another potential explanation for our results is GC-biased gene conversion (gBGC)^7^. This mutational process can temporarily increase the A/T to G/C mutation rate in small genomic regions, leading to an excess of fixed substitutions in one lineage compared to another. For example, gBGC is thought to play an important role in driving the rapid evolution of around 19% of HARs^7^. Unlike ILS, we cannot directly attribute an A/T to G/C substitution to gBGC. Therefore, we used a previously published catalog of gBGC regions in the human and chimpanzee genomes^7^. Similar to the analysis for ILS, we then estimated the proportion of the acceleration signal that could potentially be explained by gBGC.

We found evidence that a moderate proportion of HAGs and CAGs were likely influenced by gBGC (Supp. Fig. 9A-D, Supp. Tables 1, 3, and 5). However, removing all gBGC substitutions from both the human and chimpanzee lineage, we observed an equally strong pattern of substitutions occurring in more highly conserved sites in the human lineage relative to the chimpanzee lineage (p = 2.4x10^-50^, binomial test; Supp. Fig. 9F) indicating that the genome-wide bias is not explainable by gBGC. In general, whether we filter based on ILS signal, gBGC signal, or both the ratio of human- to chimpanzee-accelerated genes, proteins, and UTRs remains largely unchanged (Supp. Fig. 9A-D), suggesting that the excess of human-acceleration we observe is not driven by these processes.

**Regional mutation rate and evolutionary constraint variation beyond gBGC**

Regional variation in mutation rate (differences in mutation rate across the genome) or in evolutionary constraint can be major confounds in methods that seek to detect acceleration relative to some estimated neutral background rate. For example, if mutation hot spots are ~1 kb but the local neutral rate is estimated from 100 kb, the local rate is not an accurate reflection of neutral evolution and can lead to false positives. This is one of the issues that FASTER seeks to address. As we are comparing the relative evolutionary rate in the human and chimpanzee lineages for the same genomic regions, there can be no confounding by misestimation of neutral rates in other parts of the genome, conserved (i.e. shared by humans and chimpanzees) differences in local evolutionary constraint across the genome, or conserved differences in local mutation rate. This also makes FASTER robust to misestimation of the neutral mutation rate used in the computation of PhyloP scores as, even if this is inaccurate, this inaccuracy would essentially cancel out when comparing the human and chimpanzee lineages. Therefore, FASTER is broadly robust to these potential confounders.

The only way this can impact our results is if a genomic region has divergent mutation rate/relaxation of constraint in humans relative to chimpanzees. As we discussed relaxation of constraint in detail above, we focus here on divergence in mutation rate. As there is limited information on evolutionary divergence in mutation rate between humans and chimpanzees outside of gBGC, this could in principle potentially explain some of our results. However, this is unlikely to explain weight-driven acceleration (i.e. where a shift toward substitutions in higher PhyloP scores or larger predicted effects on CA) since this would require mutation rates to increase specifically in conserved genomic sites. As many of our results are driven by shifts toward substitutions in higher PhyloP sites or that have larger predicted effects on chromatin accessibility, this suggests that changes in mutation rate variation are unlikely to explain much of the signal we observe. In contrast, our count-driven results could more easily be affected lineage-specific changes in local mutation rate, a caveat shared by other count-based accelerated regions including HARs and HAQERs^8,9^.

**Alignment quality**

Alignment errors can, in principle, lead to signals of accelerated evolution using any methodology. Although we took extensive precautions against this (Methods), we cannot fully rule out potential alignment errors. In addition to these errors, we manually checked the alignments for all substitutions in FDR-significant human accelerated proteins and over 500 non-coding substitutions, consistently finding no evidence for alignment error. Therefore, we think it is unlikely that alignment errors have had a major impact on our results.

**Biases in ChromBPNet predictions**

In principle, ChromBPNet predictions could be biased to predict weaker or stronger effects for substitutions from the human allele relative to the chimpanzee allele or be more accurate for one of the alleles. Below, we discuss the potential impact of this on our results.

First, for the results in Figure 4 (focused on the identification of CA-HAGs and CA-CAGs), our method controls for global differences in the distribution of predicted absolute log fold-changes in CA. As a result, even if the predicted effects of the chimpanzee-derived alleles were systematically larger or smaller than those for the human-derived alleles, our method would not be biased by this. This is supported by the similar number of CA-HAGs and CA-CAGs we identify (142 and 143 respectively). In addition, the global distribution of predicted effects of chimp-derived and human-derived substitutions are highly similar both for all substitutions (median predicted absolute log fold-change of 0.021 for human-derived and 0.020 for chimp-derived) and those in sites with PhyloP > 1 (median predicted absolute log fold-change of 0.022 for both). Overall, this suggests that systematic biases are unlikely to explain the results in Figure 4.

The other result that could be affected is our finding that substitutions in conserved sites are predicted to predominantly decrease accessibility. If the model were to erroneously predict systematically lower accessibility for the ancestral allele than the human-derived allele, then this bias could produce our observed result. However, this is unlikely to be the case for two reasons. First, the fraction of substitutions predicted to decrease CA increases as sites become more conserved (Supp. Fig. 8A). Under a model of a bias toward lower CA for the ancestral allele explaining our findings, we would not expect to see this dependence as ChromBPNet does not take into account conservation. Second, we observed a highly similar pattern when using chimp-derived substitutions (Supp. Fig. 10). This is important because the model has “seen” the ancestral allele for chimp-derived substitutions but has not seen the derived allele, the opposite situation for the human-derived sites. If an erroneous bias were to exist, a likely potential cause would be that the model would be biased toward predicting that sequences that appear in some form in its training data have higher accessibility than those that did not appear in its training data. The observed symmetry between the human-derived and chimpanzee-derived substitutions in this analysis suggests this is not the case. Collectively, this suggests that whatever biases may be present in the ChromBPNet predictions, they are unlikely to explain the observations in our study.

**Distinguishing chimpanzee deceleration from human acceleration and *vice versa***

Throughout the main text, we refer to relatively faster evolution in one lineage as acceleration. However, it is possible that relatively faster evolution in one lineage could be the result of deceleration in the other lineage. To explore this possibility, we identified gorilla-derived substitutions, computed the weighted sum of non-negative PhyloP scores, normalized for the longer branch length leading to the gorilla lineage, and then computed the difference in weighted sums between human and gorilla as well as human and chimpanzee (Methods). With these data in hand, we then estimated the fraction of the signal from FASTER we observed that was due to acceleration or deceleration in one lineage relative to both other lineages. For example, *ESR1* had zero gorilla-derived nonsynonymous substitutions so had an acceleration fraction of one and a deceleration fraction of zero. Consistent with our initial assumption, many more genes had acceleration fractions of one than had fractions of zero across protein-coding regions, UTRs, and non-coding DNA (Supp. Fig. 11). For example, the mean acceleration fraction for HAPs is 0.81, indicating that only 19% of the signal of faster human lineage evolution was the result of deceleration in the chimpanzee lineage relative to gorilla (information on the gorilla lineage can be found in Supp. Tables 1, 3, and 5). Overall, we interpret this as evidence the bulk of the signal we observe can be attributed to lineage-specific acceleration rather than deceleration.
