## Supplemental Text 2 for "Adaptive loss of function accelerated the evolution of ancient and modern human cognition"

Although our results are consistent with previous evidence of human-biased accelerated evolution in non-coding regions, our finding of more HAPs than CAPs might appear to contradict previous reports of more positive selection in protein-coding regions in the chimpanzee lineage^1–3^. However, methodological differences likely explain this apparent discrepancy. Methods used in those studies, like dN/dS and Ka/Ks, represent an “average” of both positive and negative selection, biasing them toward detecting positive selection only in genes that also lack strong negative selection—most often immune and reproductive genes. In contrast, our approach gives greater weight to more conserved positions, and thus tends to have greater power in more conserved proteins. Indeed, although chimpanzees have 4% more nonsynonymous substitutions at less conserved (PhyloP < 3) sites, this trend reverses at conserved (PhyloP > 3) sites, with humans having 10% more substitutions (data not shown). As 80% of human/chimpanzee nonsynonymous substitutions have PhyloP < 3, these changes in faster evolving sites dominate the results of any methods that treat all nonsynonymous changes equally, including dN/dS and Ka/Ks. We propose that although the human lineage has slightly lower global Ka/Ks than chimpanzee, the human amino acid changes have likely had greater functional impact on processes that are generally highly conserved across mammals, such as nervous system or heart development.

1.	Arbiza, L., Dopazo, J. & Dopazo, H. Positive Selection, Relaxation, and Acceleration in the Evolution of the Human and Chimp Genome. *PLoS Comput. Biol.* **2**, e38 (2006).

2.	Bakewell, M. A., Shi, P. & Zhang, J. More genes underwent positive selection in chimpanzee evolution than in human evolution. *Proc. Natl. Acad. Sci.* **104**, 7489–7494 (2007).

3.	Vamathevan, J. J. *et al.* The role of positive selection in determining the molecular cause of species differences in disease. *BMC Evol. Biol.* **8**, 273 (2008).
