## Supplemental Text 3 for "Adaptive loss of function accelerated the evolution of ancient and modern human cognition"

**Correcting for global shifts in the conservation of divergent sites**

There are two potential explanations for why there is more acceleration in the human lineage. First, there may simply be a greater number of substitutions in conserved sites genome-wide in the human lineage, possibly due to genome-wide decreases in the efficacy of negative selection in humans. Second, the substitutions in conserved sites might be more clustered in specific genomic regions in the human lineage. To explore the first explanation, we computed the ratio of human-derived substitutions to chimpanzee-derived substitutions genome-wide in increasingly strict PhyloP bins. Notably, there was an increasing bias toward a greater number of human-derived substitutions as the conservation of the sites increased, reaching approximately 20% more in the human lineage in the most conserved bins (Supp. Fig. 2A-B, p = 6.9x10^-67^ for PhyloP > 6). This at least partially explains findings of greater acceleration in the human lineage.

However, there may also be greater clustering of substitutions in conserved sites near specific genes and CREs in the human lineage. If this were the case, it would suggest that human acceleration disproportionately affected specific genes and pathways and possibly provide clues as to the genetic basis of uniquely human complex traits. Exploring this requires controlling for the global shift toward higher PhyloP for human-derived substitutions. While this would be difficult to accomplish using previously published methods, our method can straightforwardly do so for substitutions in non-coding and untranslated regions (but not protein-coding changes). To do this, we bin substitutions by PhyloP score and then assign substitutions to the human or chimpanzee lineage within each bin. For example, if 60% of substitutions with PhyloP > 10 occurred in the human lineage, then there would be a 60% probability of assigning a substitution with PhyloP > 10 to the human lineage during the sampling procedure. To focus on the possibility that human-derived substitutions are more clustered in specific genomic regions, we use this strategy for all analyses related to UTRs or non-coding DNA. Because this strategy removes the signal of genome-wide human acceleration, it makes our results an underestimate of the actual extent of human acceleration. In addition, for all analyses related to UTRs or non-coding DNA, we incorporated an approach that controls for mutation rate in every trinucleotide context as well as differences in the distributions of trinucleotides for either UTRs or non-coding DNA near each gene (see Methods for implementation details).
