## Supplemental Text 4 for "Adaptive loss of function accelerated the evolution of ancient and modern human cognition"

We next explored the relationship between conservation and recent positive selection on nonsynonymous substitutions. We have previously hypothesized that selection in protein-coding regions in humans was a mix of positive selection in highly conserved sites that is human-specific and in rapidly evolving sites that is broadly shared across mammals^1^. Indeed, this is apparent as a greater spread in the PhyloP distribution for positively selected sites relative to unselected sites (Supp. Fig. 13A). To disentangle these two classes of selected sites, we stratified them by strength of selection, reasoning that sites under stronger selection would be more likely to be recurrently positively selected during mammalian evolution. Surprisingly, we found the opposite: sites with weaker (but still significant) positive selection had generally more negative PhyloP scores than unselected sites (Supp. Fig. 13B, p = 0.058, Mann-Whitney U test), whereas those undergoing stronger selection had more positive PhyloP scores (Supp. Fig. 13C, p = 0.023, Mann-Whitney U test), suggesting that human-specific or nearly human-specific nonsynonymous substitutions are under particularly strong positive selection during recent human evolution. In contrast, there was no significant difference in the distributions for negatively selected derived substitutions (Supp. Fig. 13D-F).

1.	Starr, A. L., Palmer, M. E., Gao, J., Nichols, C. G. & Fraser, H. B. A framework to detect positive selection using variant effect predictions reveals widespread adaptive evolution of human neurons. Preprint at https://doi.org/15.1151/2025.09.11.675696 (2025).
