## Supplemental Text 5 for "Adaptive loss of function accelerated the evolution of ancient and modern human cognition"

It is worth noting that our results on modern human evolution have several important limitations. First, we cannot determine whether the cognition-associated substitutions in conserved sites are the causal variants underlying the GWAS signal. Second, several of the GWAS traits included in our analysis (e.g. depression and educational attainment) are known to have substantial environmental confounding and phenotypic heterogeneity, such that GWAS effect estimates may partly reflect gene-environment correlations or indirect genetic effects (e.g. substitutions that alter parental behavior, indirectly affecting educational attainment) rather than direct biological effects of the variants themselves^1,2^. Third, it is unknown what traits drove the selection that acted on these variants. Many of these traits are modern constructions (e.g. educational attainment) and, even for neuropsychiatric traits that likely existed 10,000 years ago (e.g. schizophrenia), selection on linked phenotypes unrelated to cognition could explain the signal for positive selection. Finally, whether more ancient selection impacted cognition to the same extent remains to be determined.

1.	Abdellaoui, A., Dolan, C. V., Verweij, K. J. H. & Nivard, M. G. Gene–environment correlations across geographic regions affect genome-wide association studies. *Nat Genet* **54**, 1345–1354 (2022).

2.	MDD Working Group of the Psychiatric Genomics Consortium *et al.* Minimal phenotyping yields genome-wide association signals of low specificity for major depression. *Nat Genet* **52**, 437–447 (2020).
